# Beyond Locomotion: How Specialized Motor Rhythms Enable Vertebrate Escape from Capture

**DOI:** 10.1101/2025.09.08.674955

**Authors:** Saeed Farjami, Andrey Palyanov, Hong-Yan Zhang, Valentina Saccomanno, Robert Merrison-Hort, Andrea Ferrario, Roman Borisyuk, Joel Tabak, Wen-Chang Li

## Abstract

Escape behaviors following capture are crucial for survival, yet their underlying neurobiological mechanisms remain poorly understood. We investigated how *Xenopus laevis* tadpoles use struggling movements to escape head restraint. High-speed video tracking revealed a stereotyped sequence of body flexions with distinct kinematics during capture and release. We further recorded motoneuron and motor nerve activity along the body axis during fictive struggling to reconstruct biologically realistic struggling commands, to drive the movement of a biomechanically detailed tadpole model. Simulations showed that struggling - characterized by long-duration, low-frequency, caudorostral muscle activation - was optimized to generate escape forces. Notably, hydrodynamic thrust alone proved insufficient for release. However, direct mechanical interactions between the tadpole’s body and the restraining object generated additional reactive forces that facilitated escape. These findings demonstrate how animals use coordinated motor outputs and body mechanics to interact with environment to generate maximal freeing forces as the fundamental escape strategy.

## INTRODUCTION

Most animal movements, from locomotion to escape, rely on coordinated neuronal and muscle activation along the body axis ^1,2^. One type of coordination is the propagation of neuronal and muscle activities along the anterior-posterior or longitudinal body axis, which has been seen in the movement of many invertebrates ^3–10^, lower vertebrates ^11–17^, mammals ^18,19^ and even humans ^20,21^. In most cases, this is to coordinate locomotion like swimming, crawling or walking. It is also observed in non-locomotive movements like scratching ^22^ or struggling ^23,24^. . Unlike in locomotion ^3,11,12,15–17,24^, the functional significance of coordinated muscle contractions in non-locomotive movements like struggling post-capture remains less understood. In the case of capture, animals facing tight physical constraint often employ specialized motor patterns distinct from typical locomotion ^25,26^. These motor patterns must enable the animal to interact directly with the captor, rather than solely relying on propulsion against the environment..

When captured by a predator or trapped in a tight space (e.g. burrowed feeding), soft body animals can easily escape by elongating and thinning their body ^3,6^. Other animals employ backward locomotion if there is sufficient space ^9,27^. When the grip is tight, animals produce different types of motor pattern, like struggling in tadpoles ^28,29^ (supplementary video 1) and zebrafish ^24^, lamprey ^27,30^, salamander ^31^, newt embryos ^32^ or atypical tail flips in juvenile crayfish ^33^. In comparison with normal locomotion, struggling produces longer muscle contraction cycles with slower and reversed propagation along the body, which can help the tadpole escape (supplementary video 2). In addition, locomotion is normally triggered by a brief sensory stimulation or descending commands from brain motor control centres and, once started, locomotor activity is self-sustaining. In contrast, struggling requires continuous mechanosensory stimulation (as in the case of capture). Although the general principles of the underlying motoneuronal activity for struggling have been explored *ex vivo* in immobilized tadpoles ^23,28,34,35^, how these patterns generate effective escape movements *in vivo* has been unknown. For example, can struggling movements generate backward thrust in water so tadpoles move backwards, or help tadpoles free themselves?

To bridge the gap between motoneuronal activity and actual struggling movements, we combined high-speed video tracking of tadpole kinematics with detailed *ex vivo* electrophysiological recordings of motoneuron activity and three-dimensional (3D) biomechanical Virtual Tadpole (VT) modelling ^36^. By driving the VT model with diverse motoneuron activity patterns, including both natural and altered struggling commands, we systematically determined the contribution of specific motor rhythm parameters to the efficacy of escape. Our findings reveal that the distinctive features of the struggling rhythm - its low frequency, high spike count, and caudorostral propagation - are biomechanically essential for generating the forces required to free the tadpole from a physical grip. We demonstrate that escape from capture relies on the tadpole’s physical interaction with the gripping object, rather than solely on hydrodynamic thrust.

## RESULTS

### High-speed videos during grip and release reveal a stereotypical sequence of tadpole movements

Tadpoles produce vigorous struggling movements when gripped by a predator like a damselfly nymph (Supplementary Video 1 and 2) ^28^. Upon release, tadpoles then go through transitional contractions before swimming forward. We used the deep learning tool DeepLabCut for automatic tracking of tadpole movements ^37^. This enabled us to make detailed analyses of tadpoles’ body kinematics. We analysed high speed videos (240 frames per second) of 25 different tadpoles in response to grip and release.

Previous studies have shown that holding tadpoles by their heads or stimulating the rostral trunk of immobilised tadpoles are the most reliable way to induce struggling behaviour or fictive struggling with caudorostrally propagating rhythms ^23,28,34,35^. Directly holding the miniscule tadpole easily crushes the animal. To simulate the capture of tadpoles by predators and evoke struggling reliably, two fine steel pins were inserted side by side ∼ 1.5 mm apart in the bottom of a sylgard-lined petri dish (Fig.1A_1_). The tadpole was then placed dorsal-side-up between the pins with its rostral trunk supported by the pins. A pair of hand-held forceps were then used to gradually close the gap between the pins, applying pressure to grip the tadpole until it started to produce coiling and struggling movements. Gripping by the pins/forceps was released after a few seconds. The pins then sprung back to open the gap. Tadpoles typically backed out of the gap, produced coiling multiple times and swam away (Supplementary video 3).

From the videos, we identified 4 types of movement occurring in a fixed sequence: (i) initial coiling at onset of grip in which the tadpole took a C- or O-shape (once, to one side, in 17 videos and twice, to opposite sides, in the other 8 videos); (ii) struggling with the tadpole body taking S-shapes; (iii) transitional coiling after grip release during which tadpoles took an O-shape in most cycles; (iv) swimming with smaller body bends in S-shapes (Fig.1A_2_). To analyse the movements we used DeepLabCut. Eleven evenly distributed points along the tadpole longitudinal body axis were marked at the training stage and tracked. We calculated 9 body curvature angles in radians from the coordinates of the 11 points returned after tracking (Fig.1A_3_). All 9 angles were added to generate an additional trace (sum of angles) indicating overall body shape. In close to circular body shapes (C-or O-shape), the sum of angles peaks at close to 2 Pi radians, which is higher than the S-shapes typical of swimming and struggling (Fig.1B).

**Figure 1.**
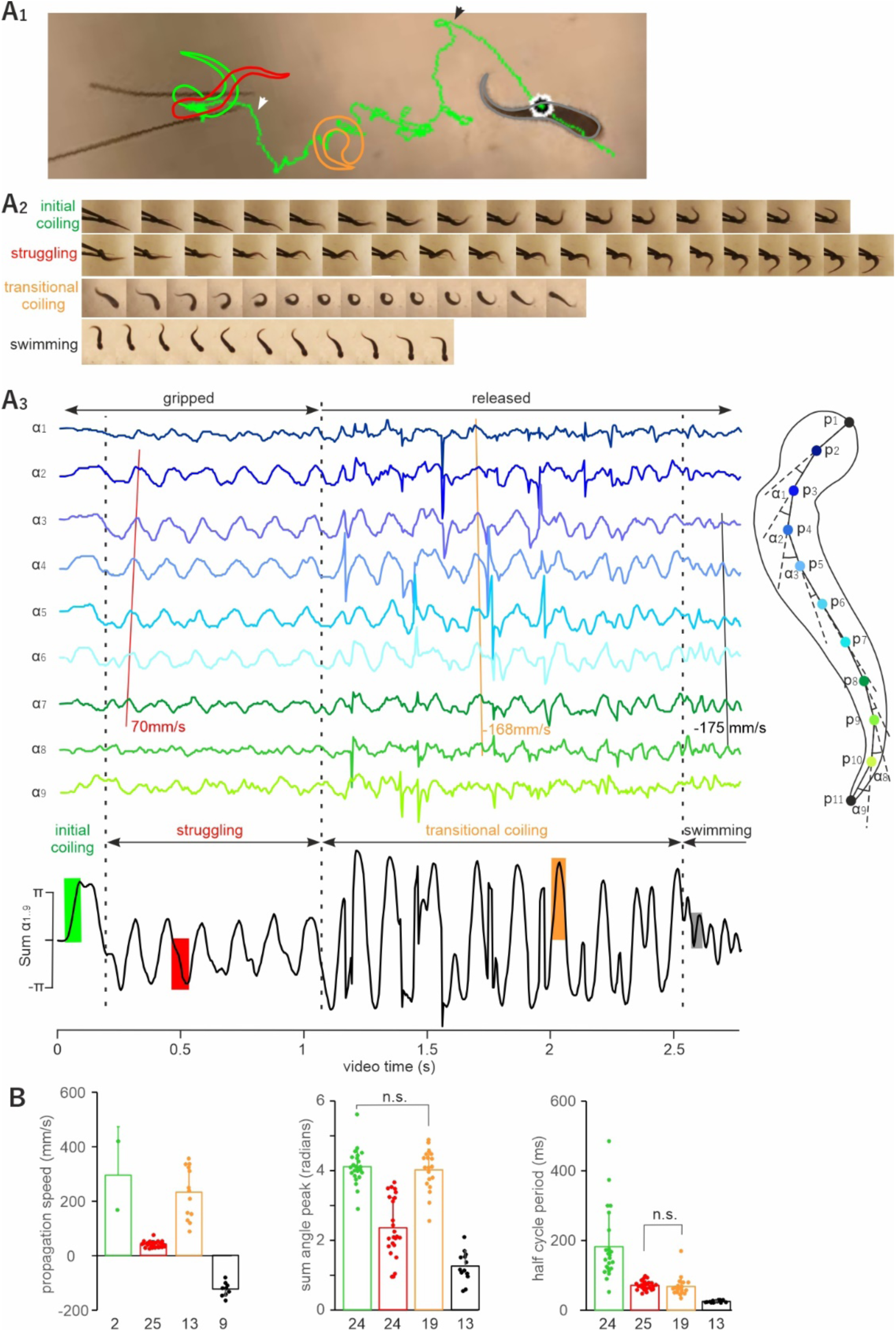
*DeepLabCut video tracking identifies four types of movement following gripping the tadpole rostral trunk and its release. **A******1****. Tadpole mass centre track (vertex near P4 on A3 diagram on the right, green) for the video analysed in **A******2-3*** *with body shapes showing initial coiling (green), struggling (red), transitional coiling (orange) and swimming (grey). White and black arrowheads indicate points of movement transition from struggling to coiling and coiling to swimming, respectively, corresponding to the middle and right vertical dashed lines in **A******3****. **A******2*** *Consecutive frames of high speed videos show tadpole body shape during each type of movement (colored/shaded periods on the Sum α1-9 trace in **A******3****). **A******3****. Nine tadpole body curvature angles calculated using the coordinates of the 11 points along the longitudinal axis (p1-p11, left traces colour-matched with dots on the righthand tadpole diagram). Solid straight lines (red, yellow, and black) show regression lines of angle peaks for estimating propagation speed of curvature changes. The black trace below shows the sum of all 9 angles with different types of movement annotated (α1-9). **B**. Comparison of propagation speeds, peaks in the sum of angles and half cycle periods for the four types of movement (bars and symbols are colour-matched with annotations in **A******1-3****). The tadpole body length is about 5mm in **A**. One-way ANOVA confirms the differences in all three measurements (all pair-wise t-tests with significance at p<0.001 except for the two pairs labelled with n.s.; for propagation, initial coiling measurements and 4 transitional coiling measurements with “infinite” speeds were excluded in ANOVA). Sample sizes are given beneath bars*.

Using the angle traces generated by DeepLabCut, we measured the average propagation speed from the regression of angle peaks or troughs, the sum of angles peak (given as half of peak/trough difference), and the duration of individual bends (to one side only, half cycle period) (Fig. 1A). One-way ANOVA identified differences between all three measurements across the four types of movement (Fig.1B, *p* < 0.001). The initial coiling differed from struggling in that its contraction half cycle was longer (183 ± 100 vs 71 ± 12.6 ms) with higher peaks in sum of angles (4.1 ± 0.5 vs 2.4 ± 0.9 Rad, both *t*-test, *p* < 0.001, Supplementary video 3). Body curvature angles, however, did not have clear peaks or troughs during the initial coiling, making it hard to estimate the propagation speed in 23 videos. In the remaining two videos, propagation speeds of 168 and 420 mm/s were estimated, much higher than the average propagation speed for struggling (41.5 ± 11.4 mm/s) but similar to the speed for transitional coiling (232 ± 95 mm/s excluding 4 with propagation delays too small to be estimated), suggesting the near-synchronous nature of muscle contractions in both types of coiling. The sum angle peaks in both types of coiling were also similar (4.1 ± 0.5 vs 4 ± 0.6 Rad, *t*-test, *p* > 0.05) and higher than those for swimming and struggling. The direction of propagation was caudorostral (positive values) for both types of coiling and for struggling. The half cycle periods for struggling and transitional coiling were comparable (*t*-test, *p* > 0.05). Moreover, the duration of transitional coiling was not correlated to the duration of struggling (𝑟^!^= 0.0002). Swimming had shorter half-cycle periods and lower peaks in the sum of angles than the other three movement types and swimming curvature waves propagated rostrocaudally (negative values, all *t*-test, *p* < 0.001, Fig. 1B).

Previous high-speed video analyses revealed that during struggling the bends originated at approximately 2/3 of the tadpole body length from the head and propagated from there towards both the head and the tail ^28^. We identified similar curvature starting points at 70 ±8.2% of tadpole lengths from the front of the head in 16 of the 25 videos, whereas no clear propagation towards the tail could be detected in the other remaining 9 videos.

### Motoneuron activity during fictive struggling and swimming reveals the same stereotypical pattern as body movements

Green and Soffe ^28^ categorised the motoneuron activity underlying struggling, transitional coiling and swimming. To define motoneuron activities underlying initial coiling, we made simultaneous recordings from spinal motor ventral root (v.r.) using two or three suction electrodes at muscle clefts between the 2^nd^ and 15^th^ myotomes in immobilised tadpoles (Fig. 2**A****_1_**). To evoke fictive coiling and struggling we either used mechanical stimulation to the rostral trunk skin or applied electrical current pulses (1ms in duration at 30 Hz). This allowed precise measurement of burst duration.

**Figure 2.**
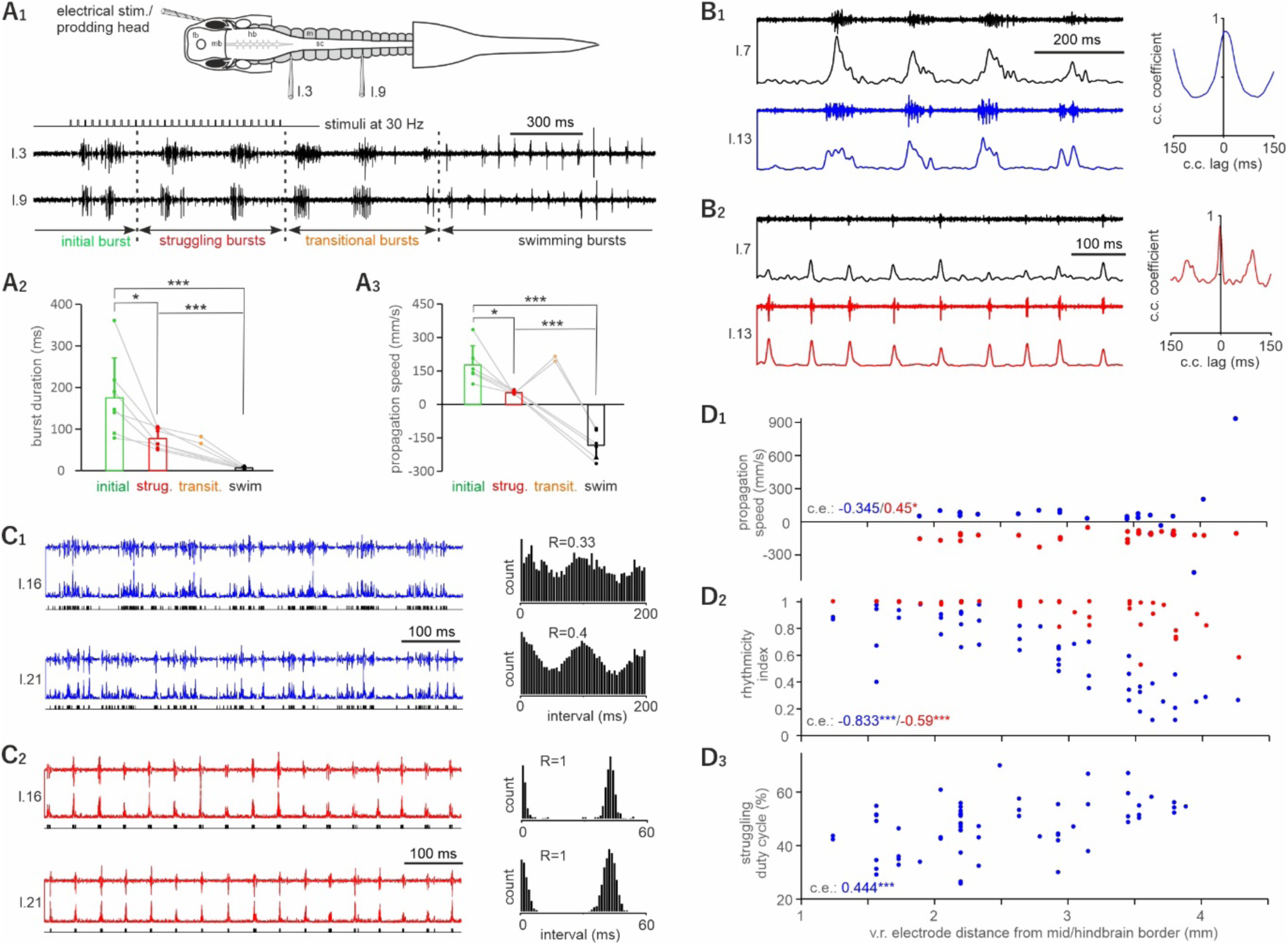
*Ventral root (v.r.) recordings identify four types of bursting activity in response to head stimulation in immobilised tadpoles. **A**. Comparing initial, struggling, transitional and swimming bursts. The diagram in **A******1*** *shows one experimental setup where two v.r. recordings are made simultaneously opposite to the stimulated, right side (l.3 means myotome cleft between the 3^rd^ and 4^th^ myotomes posterior to the otic capsule on the left side). Examples of v.r. recordings following repetitive electrical stimulation to the tadpole right head skin (**A******1****). Arrowed lines between vertical dashed lines show periods with initial, struggling, transitional and swimming bursts (color-coded as in* Fig. 1*). **A******2*** *and **A******3*** *compare burst duration and propagation speed of the four types of bursts (all two-tailed paired t-test, n= 7 tadpoles). **B** Struggling (**B******1****) and swimming (**B******2****) activities in two simultaneous channels on the left side (l.7 and l.13) rectified and smoothed for cross-correlation analyses, the peak lag of which (close to 0ms) is used for calculating rhythm propagation speed. **C** Triggering events by struggling (**C******1****) and swimming (**C******2****) activities in two simultaneous channels on the left side (l.16 and l.21) after rectification. Event interval distributions are shown on the right for calculating the rhythmicity index (R, see methods for details). **D******1*** *-**D******3*** *Correlating propagation speed, rhythmicity indexes (struggling: blue dots; swimming: red dots) and struggling duty cycles with v.r. electrode locations (Pearson correlation for rhythmicity, Spearman’s rank correlations for others). * and *** denote p<0.05 and p<0.001, respectively*.

In 17 out of 21 trials with clear struggling outputs evoked (*n* = 7 tadpoles, 1-5 trials each), there was one initial long v.r. burst (Fig. 2A_1_), which occurred on the side opposite to stimulation (16 out of 17 trials). This long initial burst was followed by rhythmic bursts characteristic of struggling. The gap between the initial burst and subsequent struggling bursts on the same side was 252.4 ± 100.6 ms. The duration of the initial burst (174.8 ± 95.8 ms) was longer than the duration of struggling bursts (76.9 ± 23.9 ms, *p* < 0.05, *n* = 7 tadpoles, paired student t-test, Fig. 2A_2_). The longitudinal propagation speed of the initial burst (178.8 ± 85 mm/s) was significantly higher than that for struggling bursts (53.8 ± 8.1 mm/s, paired t-test, *n* = 6, *p* < 0.05, Fig. 2A_3_). This long and fast propagating initial burst likely commands the initial coiling exhibited at the start of gripping in high-speed videos (Fig. 1).

After the stimulation stopped, in 2 out of 7 tadpoles, transitional bursts were observed. The transitional burst duration was in the range of struggling and initial bursts but the propagation speed of transitional bursts was only similar to that of initial bursts. Swimming burst duration was shorter than that for the other three types of bursts and the propagation was in the opposite direction, i.e. from head to tail (Fig. 2A_1-3_). The high propagation speed of transitional bursts (quasi-synchrony) reflects the transition from caudorostral propagation (struggling) to rostrocaudal propagation (swimming) ^28^.

Together, the results from Figs. 1 and 2 suggest that the initial coiling, struggling, transitional coiling, and swimming observed in high-speed videos are generated by patterns of initial bursts, struggling bursts, transitional bursts, and swimming bursts recorded from ventral roots in immobilised tadpoles.

### Differences in electrical activity patterns between rostral and caudal body locations

We next extended v.r. recordings more caudally to identify if there was any longitudinal difference in motoneuron bursting, and if there was a change of propagation direction for struggling rhythms at the two-third body length position, as seen in videos. We estimated the propagation delay, between two v.r. recordings from the same side after rectification and smoothing, as the peak cross-correlation lag time close to 0 (Fig. 2B, see methods). Propagation speeds for struggling rhythms did not change along the tadpole longitudinal body axis (*n* = 26, Spearman’s rank correlation between propagation speed and electrode position). In contrast, swimming rhythm propagation speed decreased towards the tail locations (*n* = 26, Spearman’s rank correlation, *p* < 0.05, Fig. 2D_1_).

There appeared to be more sporadic v.r. bursts towards the tail. We used a rhythmicity index to quantitatively evaluate this (Li et al., 2010, also see methods), after threshold-triggering events in the rectified v.r. recordings (Fig. 2C, black traces). There was a decrease in both struggling and swimming rhythmicity indexes towards the tail (struggling *n* = 53, swimming *n* = 47, both Pearson correlation between rhythmicity index and electrode position, *p* < 0.001). While swimming rhythmicity index was close to 1 at the majority of recording sites, the struggling rhythmicity index at the same location was always lower (*p* < 0.0001, *n* = 47, paired *t*-test, Fig. 2D_2_).

Struggling burst duration was also determined by visually grouping neighbouring events (e.g. Fig. 2C_1, 2_, event channels below v.r. traces). There was an increase in struggling duty cycle towards the tail (Spearman’s rank correlation, *n* = 61, *p* < 0.001, Fig. 2D_3_). These data show that at the CNS level there is no clear change of struggling rhythm propagation direction, contrasting with body kinematics analysis that shows bi-directional propagation from the two-third body length position. Furthermore, the struggling rhythms become less rhythmic towards the tail.

### Motoneuron distribution along the spinal cord and their spiking activity during fictive struggling

Motoneurons innervating swimming myotomes are distributed from the caudal hindbrain throughout most of the tadpole spinal cord. The staining of acetylcholine transferase (ChAT) in whole-mount tadpoles had limited success previously, probably due to the low level of antigens at early developmental stages. Here, we used a CUBIC-based clearing method ^38^ in whole-mount immunostaining to visualise potential motoneurons. The clearing step significantly improved antibody penetration through the tissue and visibility of fluorescence. This enabled us to reveal ChAT-positive cells in the caudal spinal cord beyond 3.2 mm from the mid/hindbrain border, the origin of rostrocaudal coordinates (0 mm), where few motoneurons were successfully labelled through HRP backfilling (Fig.3)^39^. The average number of ChAT-positive cells on each side of the caudal hindbrain and spinal cord was 289, higher than the total of 248 motoneurons revealed in backfilling experiments (*n* = 5 tadpoles, Fig.3A).

**Figure 3.**
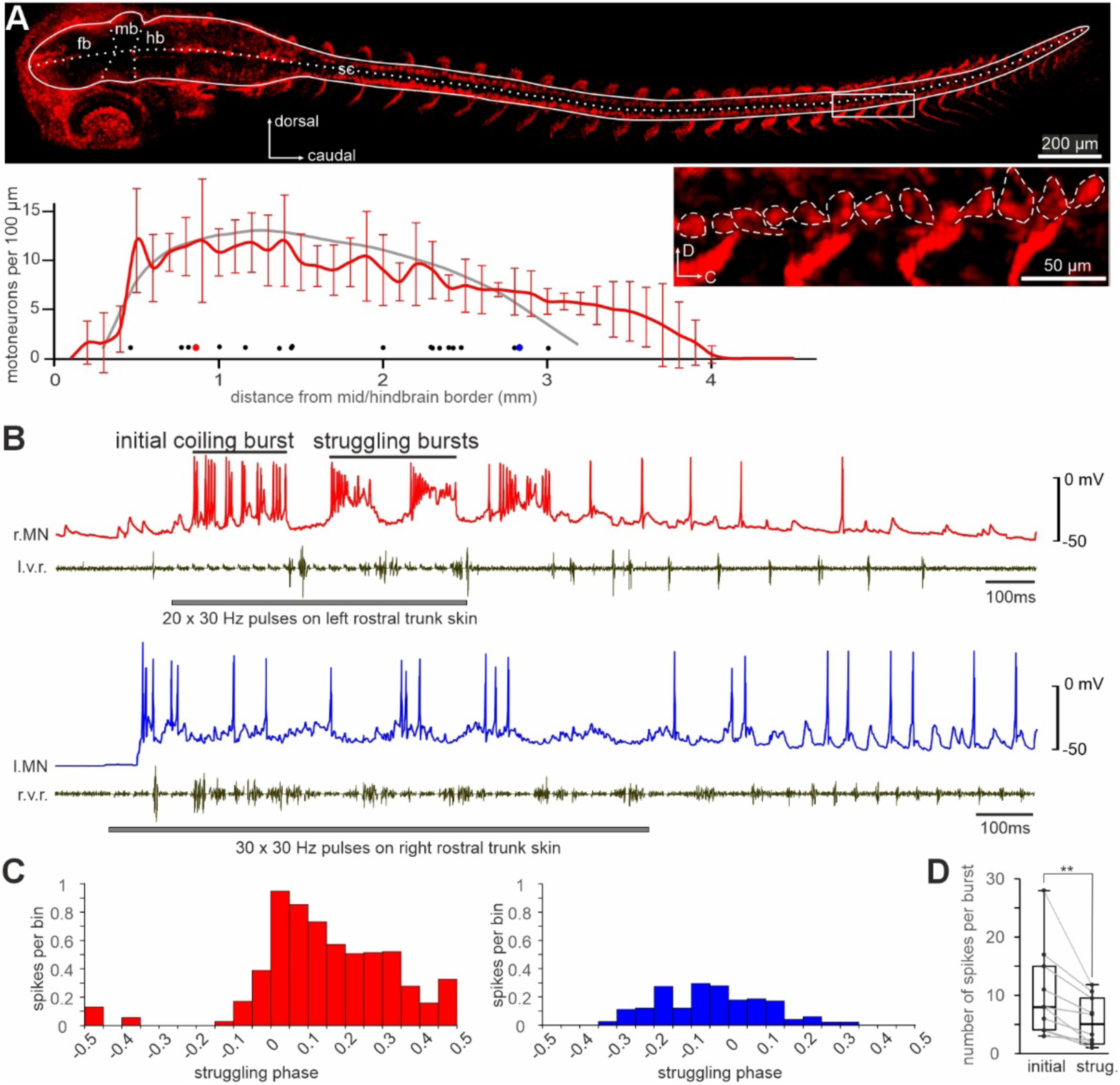
*ChAT staining of tadpole CNS neurons and motoneuron spiking during fictive initial and struggling bursts. **A**. Whole-mount immunostaining labelling of ChAT-positive cells (tilted dorsal view). White solid line delineates the CNS profile. Dotted lines mark the midline and borders between forebrain (fb), midbrain (mb) and hindbrain (hb). sc is spinal cord. Rectangular box is magnified in inset, with dashed lines circling individual motoneuron somata. Red curve indicates average motoneurons per 100 µm with SD of hindbrain and spinal cord on each side, averaged from 5 tadpoles at different longitudinal locations from mid/hindbrain border (0). Grey curve shows the number of motoneurons per 100 µm from previous backfilling experiments* ^39^*. Filled circles below curves indicate the longitudinal locations of recorded motoneurons (see B and C), with red and blue ones corresponding to red and blue traces in B. **B**. Two example motoneuron recordings following repetitive skin stimulation (horizontal grey bars) to the rostral trunk skin to evoke struggling, with simultaneous v.r. recordings (black). **C**. Phase distribution of motoneuron spikes during struggling, after caudorostral delay corrections* ^28^ *(also see methods). Red bars are for rostral motoneurons and blue bars are for caudal motoneurons. 0 means motoneuron spiking is in synchrony with ipsilateral v.r. discharge and 0.5/-0.5 means they are in anti-phase. **D**. Number of motoneuron spikes per initial and struggling burst (n = 10, two-tailed paired student t-test, ** means p < 0.01)*.

Whole-cell recordings were next made from 19 individual motoneurons from different tadpoles, located 0.46 to 3 mm from the mid/hindbrain border, to monitor their activity during fictive struggling evoked by repetitive electrical stimulation to the rostral trunk skin (Fig.3B). In trials with initial (coiling) bursts, 10 motoneurons fired 10.4 ± 7.8 spikes during the initial bursts, higher than 5.6 ± 4.1 spikes per struggling cycle (soma locations: 0.46 to 2.83 mm from mid/hindbrain border, *p* < 0.01, paired t-test, Fig. 3D). During struggling, 9 motoneurons located less than 1.5 mm from the mid/hindbrain border were grouped as rostral motoneurons. They fired 6.1 ± 4.1 spikes per cycle, higher than 1.9 ± 0.9 spikes for their more caudal counterparts (*n* = 10, *p* < 0.01, independent samples t-test, Fig. 3B, C). Their spike phase distributions during struggling cycles given in Fig. 3C were produced after calibrating the cycle period-dependent rostrocaudal propagation delay for struggling bursts ^28^. Despite more vigorous spiking in rostral motoneurons during struggling, the normalised phase distributions of spikes in the rostral and caudal motoneuron groups were similar (Independent sample Mann-Whitney U test, *p* = 0.06).

### Generating motoneuron spiking commands to drive virtual tadpole (VT) movement

We recently developed a biomechanical virtual tadpole model (VT) to simulate body movements, in which motoneuron activities generated by a detailed CNS model ^36^ (also see methods) contract VT muscles to generate realistic swimming behaviour.

Here we use the VT model to bridge the gap between our experimental studies of electrical activity during fictive struggling (Fig. 2) and of kinematics of struggling movements while a tadpole is gripped by forceps (Fig 1). To do that we generate spatio-temporal patterns of motoneuronal bursting activity (“surrogate data”) which mimic the properties of ventral root recordings (burst characteristics at different rostrocaudal positions, speed and direction of propagation). These patterns innervate the muscles of the VT to generate struggling movements while the tadpole is free or gripped by virtual forceps. The struggling behaviour of VT appears realistic (e.g. supplementary video 4) and our analysis of VT struggling (below) demonstrates characteristics of tadpole movements. Simulations of the VT model enabled us to determine how different burst parameters (duration, speed and direction of propagation) affected the VT’s movements and its ability to escape from virtual forceps.

We first expanded the distribution of motoneurons in VT to include more caudal spinal cord. Our previous VT model only had motoneurons in the CNS region 0.5 – 3.0 mm from the mid/hindbrain border ^36,40^, based on backfill experiments (^39^, Fig.3A, grey line). Our new ChAT staining revealed positive cells in the spinal cord beyond 3.2 mm from the mid/hindbrain border (Fig.3A, red line) whilst v.r. discharges were recorded near the end of the tadpole tail (Fig.2D). Although some motoneurons have central axons running caudally in the spinal cord before exiting to innervate myotomes, data on the axonal projection of such motoneurons is lacking. We simply extended the distribution of motoneurons from the previous rostrocaudal (R-C) coordinates of 0.5 – 3.0 mm to 0.5 – 4.8 mm from mid/hindbrain border, increasing the number of motoneurons on each side from 248 to 311 (Fig. 4A_2_, black line).

**Figure 4.**
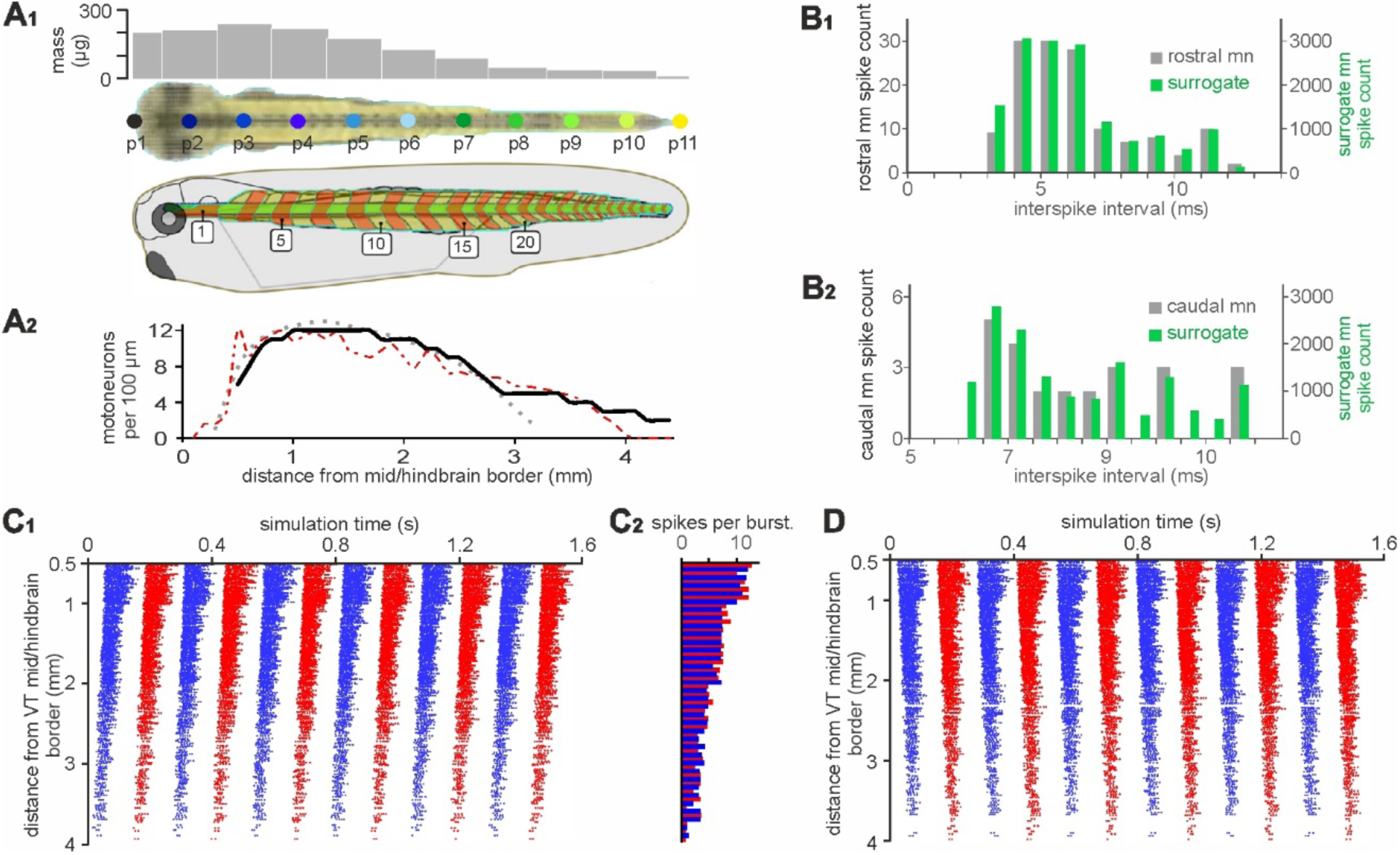
*Generating surrogate motoneuron spiking data to drive VT movement. **A******1****. VT model with muscle segments (middle: dorsal view; bottom: side view) and mass distribution at body segments around the 11 evenly separated longitudinal body axis points p1-11 (top grey bars). Numerals inside white boxes on the side view drawing indicate swimming myotome numbering. **A******2****. Distribution of motoneuron somata along the longitudinal axis of the CNS in VT (black solid line). Grey and red dotted lines are from backfills* ^39^ *and ChAT staining, respectively (also see* Fig. 3A*). **B******1****-**B******2****. Matching surrogate motoneuron inter-spike intervals with those in whole-cell recordings of rostral and caudal motoneurons. **C**. Surrogate motoneuron spiking in the VT for struggling. Red and blue dots show the longitudinal location of every motoneuron and their spike time in the simulation on the left and right sides, respectively (**C**1, caudorostral propagation at ∼ 40 mm/s). Bars in **C**2 shows the average spike number per motoneuron per struggling burst (red is for left motoneurons and blue is for right motoneurons, averaged for cells within the same 100 µm of CNS). **D**. Surrogate motoneuron spiking data similar to C but with a reversed direction of propagation*.

We tuned the surrogate motoneuron spiking features to match those extracted from fictive struggling rhythms (Fig. 2-3), including inter-spike intervals, struggling frequency, duty cycle, propagation direction and speed. The distribution of 14382 surrogate motoneuron inter-spike intervals was similar to that for 138 motoneuron spikes in whole-cell recordings of rostral motoneurons (Independent sample Mann-Whitney U test, *p* = 0.74). Similarly, the distribution of 14832 surrogate inter-spiking intervals for more caudal motoneurons was comparable to that for 24 motoneuron spikes in whole-cell recordings (Independent sample Mann-Whitney U test, *p* = 0.58, Fig. 4B**_1-2_**). The mean number of spikes per burst in different parts of the spinal cord are shown in Fig. 4**C****_2_**, with rostral motoneurons firing more spikes per struggling cycle than their caudal counterparts (c.f. Fig. 3C). The surrogate struggling command frequency was 4 Hz, with a duty cycle of ∼ 0.4 for the rostral VT CNS (c.f. Fig. 2D_3_), which activate the rostral myotomes for the VT to interact with the virtual forceps tips. The caudorostral propagation of surrogate motoneuron bursting was adjusted to ∼ 40 mm/s (Fig.4C), similar to that measured during fictive struggling (Fig.2A, B). To simulate the initial bursts, we fixed the number of spikes at 15 with little longitudinal propagation delays. We also computed a surrogate reverse struggling command that had the same motoneuron spiking parameters as normal struggling but with a rostrocaudal direction of propagation (Fig. 4D).

### Struggling movements can free VT from gripping forceps

As shown in high speed videos and v.r. recordings, the generation of struggling rhythms depend on continuous stimulation to the skin. When the stimulation stops, the tadpole movements transit to coiling before swimming starts.

A priori, it is not clear if the tadpoole struggling movements can move the body backwards. Therefore, we imposed surrogate struggling commands (Fig.4C_1_) on the VT in free water to see if struggling movement alone could lead to backward translation (supplementary video 5). Simulation results show the VT generated curvatures with synchronous bending over the body length (Fig. 5A). Peaks of sum of curvature angles (Fig. 5C_1_, left panel) were about 4.46 ± 0.53 Rads, typical of transitional coiling (Fig.1B). When the VT was not obstructed by the water tank, it moved *forward* at a speed of 9.6 mm/s, about 4 times slower than its swimming speed (Fig. 5A). Thus, struggling motoneuron commands are ineffective at moving the free virtual tadpole backward.

**Figure 5.**
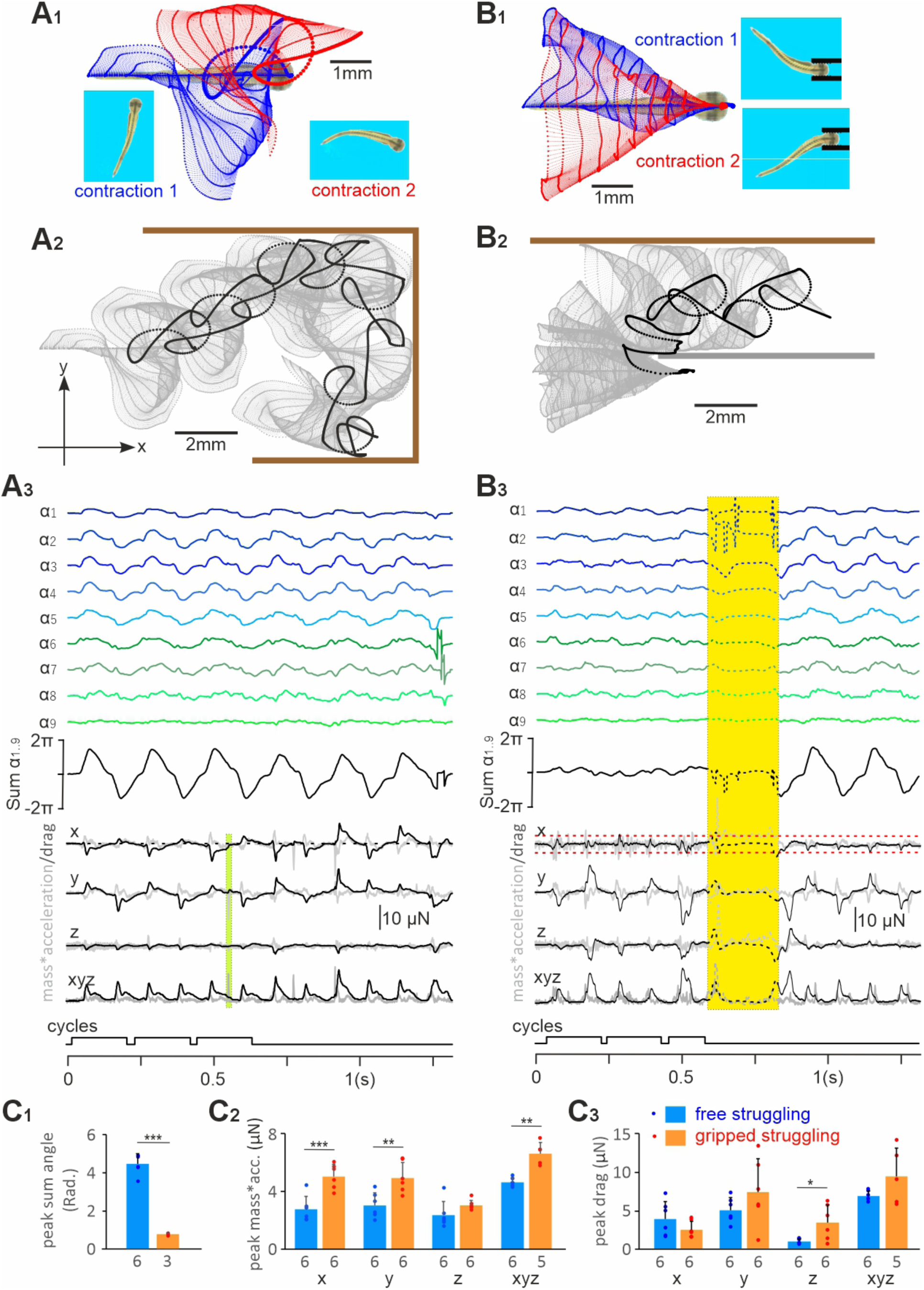
*Struggling movements do not move the free VT backwards, but can make it escape backwards from grips. **A**. Tracking the free VT struggling movement in water. **B**. Tracking struggling movement when VT is gripped by forceps at the threshold grip strength. **A******1, B1****. The movement trajectories seen from above of p1-p11 for the first (blue symbols, lines) and second (red symbols, lines) contractions with larger dots indicating p1. Screen shots show curvatures at end of contractions. **A******2, B2****. VT movement trajectories for total simulation time (grey dots and dashed lines). Large black dots are for tracking point p1. Thick brown bars indicate virtual water tank walls and thick grey bar indicates location of one side of the virtual forceps that VT touches during movement. Inset axes for drag and acceleration: x is the direction of VT longitudinal body axis at rest (positive value means forces propelling VT forward, i.e. to the right in panel A); y is the up/down direction in panel A; z is perpendicular to the xy plane. **A******3, B3****. VT body curvature angles, mass*acceleration (grey traces) and drag force. We show projections onto the three axes and the magnitude of xyz 3D vectors. “Cycles” trace indicates periods of analyses. Yellow-shaded period with dashed traces have missing tracking points due to untraceable VT orientation around moment of escape (see supplementary video 4). Green shading in **A******3*** *indicates the time when VT briefly touches the tank wall, which is excluded from the analyses. Red dashed line in **B******3*** *indicates the magnitude of the friction force from the forceps. The drag force (black curve) never exceeds this value. Surrogate motoneuron spiking data are from* Fig. 4C*. **C**. Comparing peaks of sum of angles (**C******1****), mass*acc.(**C******2****) and peak drags (**C******3****) generated by free VT struggling and gripped struggling (two-tailed independent sample t-test, *** indicates p < 0.001, ** indicates p < 0.01, * indicates p < 0.05). Numerals beneath bars are numbers of contractions/cycles*.

We lacked biologicl data to simulate a virtual predator like the damselfly larva and the flexible insect pins inserted in elastic sylgard also added difficulty to simulations. To simulate the predatory capture, we simply introduced a pair of rigid virtual forceps (see methods for details) to grip the VT head (by ∼ 1.4 mm), with the rest of the VT body length outside the forceps (Fig.5B_1_). The gripping strength was proportional to the amount of compression (represented as proportion of VT head width/h.w.) applied to the VT head by the forceps tips, directly affecting the friction between VT and the forceps (supplementary Fig.1, also see Methods). When the VT head was compressed by 0.13 times h.w., struggling movements generated by the VT did not lead to escape. When the grip was reduced to 0.124 times h.w., the VT could free itself after six struggling contractions (Fig.5B, supplementary video 4), which we considered as the maximal grip strength that VT could escape from (threhsold grip). As the grip was further relaxed to 0.118 times h.w., the VT could free itself with just one struggling contraction. We also tested surrogate commands with two initial coiling bursts. The coiling contractions were ineffective at freeing the VT, which could only escape after the first struggling contraction took place.

How were struggling contractions able to free the tadpole from the forceps grip? First, we noted that the grip restricted VT movements, leading to significant reduction of peak of sum of angles from 4.46 ± 0.53 to 0.76 ± 0.07 Rads (two-tailed independent sample *t*-test, *p* < 0.001, Fig5C_1_). We also calculated the acceleration (multiplied by the mass for comparison with drag force) and the pressure drag force in three dimensions: axis x refers to the initial VT rostral/caudal direction (positive means forward), y is perpendicular to x and z is VT dorsal/ventral direction, whereas xyz is the magnitude of the 3D vector combined from x, y and z. The peak of mass*acc was 5.05 ± 0.89 µN (x axis projection). During these struggling contractions, the water resistance to the VT body movements generates a drag force in the opposite direction to the movements. At times during a cycle, this drag force points away from the forceps, opposing the friction force, and possibly contributing to free the tadpole. However, the absolute peak/trough of the drag force in the x direction, was 2.43 ± 1.1 µN (Fig.5C_2-3_), lower than the friction force that we calculated to be 3.2 µN (see Suppl. Fig. 1). This shows that the drag force generated against the water, alone, could not free the tadpole. Direct interaction of the VT with the forceps must have produced the (extra) force that led to the VT escape.

### Movement driven by reverse swimming and struggling commands

Having demonstrated VT struggling movements could result in the VT escaping from gripping forceps, we next asked if other forms of surrogate commands could help the VT escape from the gripping forceps.

Previously we simulated VT swimming with the biologically realistic swimming rhythms generated by a CNS swimming network ^36^ (Supplementary Fig. 3A_1_), in which motoneurons typically fired a single spike per swimming cycle with spiking propagating rostrocaudally at a speed of ∼ 130 mm/s. We first generated surrogate reverse swimming rhythms by shifting the same motoneuron spiking so that the 20 Hz rhythms propagated caudorostrally at ∼ 130mm/s (see Supplementary Fig. 3B_1_). When the virtual forceps gripping was at 0.124 times h.w., corresponding to the threshold grip strength for struggling, VT did not free itself after 17 reverse swimming contractions (Fig. 6A, supplementary video 6). In comparison with free movement driven by the reverse swimming command, gripped reverse swimming generated smaller peaks of sum of angles, larger x axis mass*acc and higher drag peaks/troughs (Supplementary Fig. 3C). Although the drag had positive peaks transiently exceeding the friction of 3.2 µN between the VT and forceps, the VT did not escape at this level of grip strength. However, the VT did free itself at the fifth contraction when the grip was gradually reduced to 0.11 times h.w. and the friction between VT and forceps was 2.77 µN. Therefore, reverse swimming could help the VT escape the forceps grip but it was not as effective as struggling.

**Figure 6.**
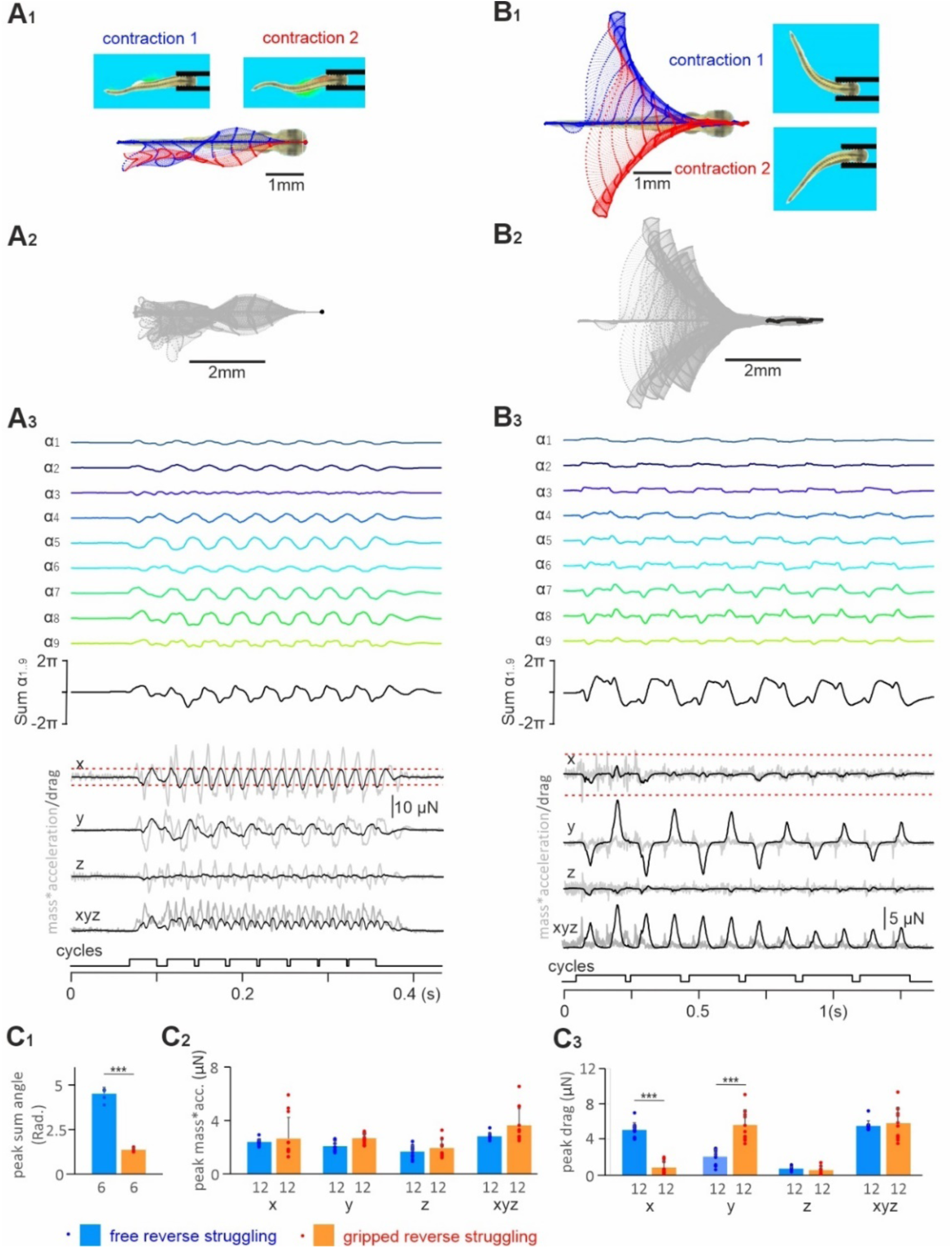
*Tracking gripped VT movement driven by reverse swimming and reverse struggling commands. **A.** VT movement driven by reverse swimming commands. **B.** VT movement driven by reverse struggling commands. **A******1-B1****. The movement trajectories of p1-11 along VT body axis of the first (blue symbols, lines) and second (red symbols, lines) contractions. Screenshots show VT body curvatures at the end of contractions. Larger dots indicate p1 trajectories. **A******2-B2****. VT movement trajectory for the total simulation time (grey dots and dashed lines). Large black dots are for p1. **A******3-B3****.VT body curvature angles, mass*acc. (grey traces) and drag (black traces) in three axes. Surrogate motoneuron spiking data in A is from Supplementary Fig3B1, and **B** is from* Fig.4D*. **C**. Comparing peak of sum of angles (**C******1****), peak mass*acc. (**C******2****) and peak/trough drag (**C******3****) generated by free reverse struggling (Supplementary Fig.2) and gripped reverse struggling (all two-tailed independent sample t-tests, *** stands for p < 0.001 and * for p < 0.05). Numerals beneath bars are numbers of contractions/cycles compared. Red dashed lines in **A******3*** *and **B******3*** *indicate friction levels between the VT and forceps*.

Secondly, we tested reverse struggling rhythms (Fig. 4D). When the VT was not held by the forceps, it turned around with each contraction (left-right yaw: 160.2 ± 17.6 degree, *n* = 12, Supplementary Fig. 2, supplementary video 7), with large peaks of sum of body curvature angles (4.39 ± 0.38 Rad, *n* = 6, Fig 6C_1_) typically seen in coiling but with a slow body curvature wave propagation speed of ∼ 90mm/s (c.f. Fig.1B). There was no consistent displacement of the VT, e.g. backward movement.

When held in forceps and the grip was stronger than 0.156 times h.w., the VT driven by reverse struggling commands did not move along the forceps axis. However, when the grip was at 0.154 times h.w. with a friction of 4.2 µN between the VT and forceps, the VT moved deeper into the forceps with each of the first four contractions (by 0.65mm) and got stuck afterwards (Fig. 6B, supplementary video 8). Peak/trough drag of 0.89 ± 0.72 µN on x axis generated by the reverse struggling rhythms in the gripped VT is lower than the friction of 4.2 µN, insufficient to move the VT inside or out of the forceps. Therefore, the movement of VT into the forceps must have been helped by the direct interaction between VT and the forceps. At gripping strengths up to 0.11 times h.w., the VT could advance about 1.43 mm into the forceps with only 44% of its length outside the forceps free to move. Consequently, the magnitude of drag generated by the VT dropped (similar to Fig. 6B_3_) and VT became stuck. When the grip strength was further decreased to 0.076 times h.w., the VT escaped from the forceps at the first contraction. These simulations show that the motoneuron bursting duration, frequency and propagation direction are all important in the VT escape and the natural struggling rhythm allows VT to escape from the strongest grip.

### Reverse swimming commands do not produce backward VT movement

Some fish species like dogfish ^41^, lampreys ^27^ eels and catfish ^16^ can reverse their normal rostrocaudal axial muscle contraction sequence in forward swimming and swim backwards, to help them back out of/into narrow crevices. Backward swimming has never been observed in tadpoles at stage 37/38 in behavioural tests. We tested whether theoretically reverse swimming commands (Supplementary Fig. 3B_1_) could allow the VT to swim backwards and generate backward thrust, potentially facilitating escapes from grips. When VT movement was driven by reverse swimming commands (supplementary video 9), the coordinates of 11 longitudinal VT axis points were tracked and body curvature angles calculated. Interestingly, the reverse swimming commands still generated rostrocaudal body curvature waves at a propagating speed of ∼ 180 mm/s (Supplementary Fig. 3B_4_), faster than 116 mm/s seen in normal swimming. Also, reverse swimming commands propelled the VT forward, at a speed of 43.15 mm/s, similar to the forward VT swimming speed of 41.16 mm/s (Supplementary Fig. 3A_2-3_, B_2-3_). Kinematics analyses showed swimming with reversed commands had larger peaks in the sum of body curvature angles (3 ± 0.14 vs 1.08 ± 0.15 Rads) and larger left-right yaw angles than those measured in normal swimming (supplementary video 10, 169.1 ± 1.1° vs 104.5 ± 7.1°, both measured when VT was free of obstacles, *p* < 0.001 and two-tailed independent sample student *t*-test). Reverse swimming commands generated larger acceleration in all three axes although the trough pressure drag force of -4.49 ± 7.02 µN on the x axis was similar to that of -3.92 ± 1.39 µN in normal swimming (Supplementary Fig. 3C). It is noteworthy that the drag remained negative when the VT was moving forward in both cases (black x axis traces in Supplementary Fig.3A_4_, B_4_), consistent with the resistive nature of drag. The simulation shows that VT is not capable of backward swimming, which could be a result of the front-heavy distribution of body mass ^16^ (Fig.4A_1_).

## Discussion

High-speed video recordings have identified the sequence of motor events following a tadpole’s head being gripped: i) initial coiling, ii) struggling; and, if the tadpole is released: iii) transitional coiling and iv) forward swimming. Electrophysiological recordings also identified four phases of electrical activity during continuous stimulation: i) long synchronous bursts, ii) bursts propagating caudorostrally; and, after stimulation is stopped: iii) transient short synchronous bursts and iv) spikes propagating rostrocaudally. Combining these data with a biomechanical model of the tadpole (Virtual Tadpole model) established a one-to-one mapping between the sequence of electrical events and the sequence of motor events. The VT model also showed that the struggling rhythm, with its slow pattern and tail-to-head coordination, is optimal to enable the tadpole to escape from grips. By comparing the friction exerted on the tadpole to the drag force generated by the water on the struggling body, we demonstrated that the force that frees the tadpole from grips results from the tadpole’s physical interaction with the gripping object.

The tadpole’s struggling motor pattern differs much from the swimming pattern in terms of rhythm frequency and coordination. This is expected, since the scenario for escaping grips is very different from that for normal locomotion. Firstly, when animals are physically restricted by captors or trapped headlong in narrow space, tactile and/or pain senses provide the main sensory inputs whereas vision or hearing may be compromised. In tadpoles, such continuous stimulation of the mechanosensory system is also essential for maintaining the struggling rhythms ^23,35^ just like scratching in turtles ^42^. Second, the purpose of escape movement is for the animal to regain freedom to move, so energy efficiency is not a priority unlike in locomotion. Third, the substratum that an animal can work with also differs. Instead of interacting with solid/semi solid surface, water or air in locomotion, the animal body is in direct contact with the gripping object. Animal escape movements must be appropriate for this specific sensorimotor context.

When a predator strikes from the head end, it makes sense for the prey to first turn around and then flee using their energy-efficient forward locomotion gait. The initial coiling we have identified resembles the turning behaviour tadpoles exhibited when suction was applied to their head ^43^, the turning response in zebrafish to strong tactile stimulation ^24^, or lamprey responding to a pinch to the head ^44^. Here, though, coiling happened after the tadpole was gripped, and our simulations show that this movement does not help free the tadpole from forceps.

As reported previously, after escaping, the transition to swimming is often preceded by a few transitional coiling bouts ^28^. The coiling movement involves near synchronous bursting of ventral roots and contraction of muscles on the same side. These transitional coiling bouts of free tadpoles do not lead to predictable, effective displacement of the tadpole ^28^. While the initial coiling may well correspond to turning, the significance of transitional coiling between struggling and swimming remains to be determined. It seems counter-productive to produce random movements after escaping the predator’s jaws, rather than quickly swimming away. Perhaps these random movements make the tadpole’s trajectory less predictable. We have previously found that following a brief stimulus a tadpole would initiate swimming after a random delay and would also start swimming away in an unpredictable direction ^45–48^. The tadpole’s vision and other sensory messages to the brain are not functional at this stage, and the tadpole does not choose an action – the action happens automatically. In the absence of decision making, the randomness provided by vigorous transitional coiling may be the best strategy to avoid recapture.

Our biomechanical model shows that struggling rhythms can help free the VT from forceps. When the tadpole is gripped, its body interacts with two substrata, water and the forceps, to generate forces. Limbless aquatic animals without pectoral fins can contract their axial muscles along their elongated body in a caudorostral order to swim backwards in water ^16,27,41,44^. However, in our simulations, driving the gripped VT using reversed swimming commands could not free the VT, unless the grip was relaxed. Moreover, it is not just the power of the struggling rhythm that frees the VT: driving the VT with a reversed struggling command only push the VT forward. Thus, we conclude that both characteristics of the struggling pattern – the stronger/slower motoneuron discharges and the caudorostral propagation – are required to free the tadpole.

What mechanisms could help the tadpole free itself by its struggling movement? First, the caudorostral struggling rhythm generates a backward pull, due to the drag force from the water opposing the lateral body movements. Second, the struggling body may interact directly with the forceps, and this produces a backward force on the body to pull it away from the grip. We have eliminated the first possibility, by calculating the drag force generated by the struggling movements. In cases of gripped VT struggling, the peak drag force opposite to the forceps axis was smaller than the friction between the forceps and VT. Therefore, there must be another force, which results from the direct interaction between VT and forceps, to free the tadpole.

How can this direct interaction between VT and forceps generate the freeing force? In our model, when the forceps are closed the VT cannot wedge them open. This may be different from real predation (e.g. capture of a tadpole by a damselfly larva) but simplifies our simulation so the VT only needs to overcome the friction force along the x axis (i.e. parallel to the forceps). From this assumption, and after carefully examining videos of the VT freeing itself, we have developed a speculative hypothesis that explains why the struggling pattern is effective at freeing the tadpole. When the VT rostral trunk muscles contract simultaneously, the rostral trunk stiffens up and works like a lever. The movement of the body on the posterior side of the lever pushes that end of the lever (mass*acc), whereas the VT contact with the supporting forceps tip functions as the fulcrum. Factors like muscle contraction pattern, forceps gripping strength/gap and the length of VT inside the forceps will affect how much force is generated at the VT headend, which could move VT either out of or deeper into the forceps (supplementary Fig.4). For struggling and reverse swimming commands, the forces generated by VT momentum pulls the VT out of the forceps. Since struggling has longer burst duration, more muscles contracting at the same time will form a longer lever helping to generate more pulling force at the VT head. For reverse struggling, the momentum generates a pushing force that pushes VT further into the forceps. Once the VT is deeper into the forceps, its movement is more restricted by the forceps and the leverage effect decreases. This reduces the pushing force and the VT becomes stuck when this force is lower than the friction between VT and forceps (Supplementary Fig.4A-C).

In the potential scenario of capture, the direct interaction of prey body and the captor is critical for survival. A similar situation occurs when animals without limbs need to back out of tight spaces. For example, when a hagfish tries to withdraw from the small opening of a wired cage grid ^49^. Slow caudorostrally propagating muscle contractions generate much less side undulations than normal forward swimming. However, the fish still manages to free itself. In this case, the caudorostral contraction wave appears to be optimised for the fish to directly interact with the physical substrate around the opening, rather than with water, likely through similar biomechanics for the tadpole struggling movement.

Furthermore, while during fictive struggling electrical activity propagates from tail to head, in the videos of struggling tadpoles we identified a point situated at about 2/3^rd^ body length from the head, from which bending waves propagated both forward and backward. The VT model confirmed that the propagation from this 2/3^rd^ point can indeed result from an electrical activity pattern that propagates forward from the tail, suggesting the tail movement is passively imposed by the more rostral myotomes, rather than active muscle contractions. This passive movement likely does not contribute significantly to propulsion since tadpole tail shape does not influence tadpole swimming performance ^50^.

To conclude, by adopting a hybrid experimental-modelling approach, we have analysed how tadpole struggling movements induced by head-gripping could generate freeing forces. In the event of capture, the prey animal will have a better chance to escape by directly engaging with the captor’s body, rather than solely with the substratum for normal locomotion.

## Methods

We injected pairs of adult *Xenopus laevis* with human chorionic gonadotropin (HCG, Sigma, UK) to induce mating so embryos could be collected and raised at different temperatures to prolong tadpole supply. Procedures for HCG injections comply with UK Home Office regulations, though pre-feeding tadpoles are considered insentient. All experimental procedures on tadpoles were approved by the Animal Welfare Ethics Committee (AWEC) of the University of St Andrews. Damselfly larvae were purchased from Blades Biological Ltd (Cowden, Edenbridge, UK).

### Video tracking

Ultra high definition (UHD) videos at 60 fps were recorded using a Pixel 9 Pro mobile phone with 2x optical zoom to capture Damselfly larva catching tadpoles. The camera was supported by a stereomicroscope frame but could be slid to follow the larva movement. Each time one damselfly larva was placed with 5-8 tadpoles in the same opaque creamy-white dish with a diameter of 75mm. We did not quantify the videos since the number of videos was small and frame rate was low. HD videos of tadpole movement resulting from head gripping using steel insect pins and forceps were made at 240 fps using a goPro hero 11 camera mounted on a trinocular stereo microscope (Brunel Microscopes, UK). The tadpole was placed in de-chlorinated water in a petri dish (diameter 60mm) lined with sylgard at the bottom. Two insect pins were inserted about 1.5mm apart. A tadpole was sucked up using a plastic pipette and placed with its head end between the two insect pins. Then a pair of forceps were used to push the pins progressively closer to apply pressure on the tadpole neck region for up to a few seconds to evoke struggling. Both UHD and HD videos were cropped at one eighth of the original speed for visual inspection (supplementary videos 1-3) and tracking. In order to track tadpole kinematics in the high-speed videos in DeepLabCut ^37^, the tadpole longitudinal body axis was marked by 11 evenly distributed points stretching from its head extremity to the tail tip. The (𝑥, 𝑦) coordinates of these points in the two-dimensional (2D) plane were registered by DLC tracking. They are used to calculate tadpole movement speed and acceleration. Tadpole body curvature angles at the intermediate 9 points were calculated using the two lines connecting neighbouring pairs of points (Fig.1A_3_). We define two vectors 𝑎⃗ and 𝑏^.⃗^ using these points: 𝑎⃗ with a start at the previous point and the end at the anchor point and 𝑏^.⃗^ with a start at the anchor point and the end at the next point. Then, the angle is calculated in radian as 𝜃 = 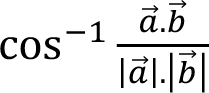 using the inner (dot) product of two arbitrary vectors. Regression was carried out to connect curvature angle peaks/troughs to estimate the propagation speed of tadpole body undulations. In the regression, we only considered angles 𝛼_2_ − 𝛼_6_ and excluded 𝛼_1_ as it did not vary much. Separate regression was conducted using 𝛼_7_, − 𝛼_9_ because in the majority of cases there was a change in propagation direction.

Also, the 9 angle measurements at each frame were added to generate a sum angle measurement. During coiling, tadpole body took a C-or O-shape, rather than an S-shape in struggling or swimming. This sum angle peak provided an effective parameter for capturing the overall body shapes. For each body bend (half cycle in case of alternating bends in struggling and swimming), we could measure the duration using the sum angle trace. At stage 37/38, tadpoles have not developed neuromuscular mechanisms to stay upright when they are moving in the water ^51^. Therefore, its body orientation became unpredictable when the tadpole was released from the mechanical grip, rendering tracking less accurate especially during the transitional coiling period. In these cases, movements were visually inspected to correct the number of coiling cycles. Due to the low frame rate, we did not calculate mass*acc and drag for these videos of real tadpole movement.

### Electrophysiology

*Xenopus* tadpoles at stage 37/38 were first anaesthetised with 0.1% MS-222 (3-aminobenzoic acid ester, Sigma, UK), then pinned onto a rubber stage so their dorsal fin could be slashed open to allow immobilisation by 10 µM α-bungarotoxin for ∼ 20 minutes. Saline ingredients included (mM): NaCl 115, KCl 3, CaCl_2_ 2, NaHCO_3_ 2.4, MgCl_2_ 1, HEPES 10, adjusted with 5M NaOH to pH 7.4. After immobilization, the tadpole was further dissected to expose muscle clefts where suction electrodes were placed to record v.r. activities. Some muscles overlaying the spinal cord were also removed to expose the spinal cord. Then a dorsal cut was made along the midline to open up the spinal cord and further cuts were made inside the neurocoel to expose neuronal somata for whole-cell recordings. The recording chamber contained a small rotatable rubber stage for mounting the tadpoles with electrically etched tungsten pins. Saline circulation was maintained at about 2 ml per minute using a peri-static pump. Patch-clamp electrodes were place onto the tadpole under an upright Nikon E600FN microscope.

Ventral root recordings were made using glass suction electrodes placed at muscle clefts at different longitudinal locations to monitor motoneuron outputs. A stimulating suction electrode was placed on the rostral trunk or head to evoke fictive struggling in immobilised tadpoles. Patch pipettes contained 0.1% neurobiotin in an intracellular solution with (in mM): 100 K-gluconate, 2 MgCl_2_, 10 EGTA, 10 HEPES, 3 Na_2_ATP and NaGTP with pH adjusted to 7.3 using KOH and had resistances ranging 10-20 MΩ. Signals were recorded with an Multiclamp 700B amplifier in current-clamp modes. Data were digitised with a CED 1401 board controlled with Signal software (CED, Cambridge, UK) with a sampling rate of 10 kHz. Stimuli to the skin were delivered through a DS-3 stimulator (Digitimer, UK), with timing controlled by Signal.

Electrophysiology recordings were analysed using Dataview (courtesy of Dr W.J. Heitler in the University of St Andrews). Threshold was set to trigger events by v.r. bursts. These events were then visually inspected and merged to form a single bursting event for each struggling cycle for calculating burst duration, struggling cycle period/frequency and duty cycle. In order to measure propagation time for struggling and swimming rhythms between different recording electrodes on the same side, v.r. recordings were rectified and smoothed, by way of moving average of a 2.5 ms data window with 4 iterations. Then the peak time close to 0 on the cross-correlation curve was determined as the propagation delay. Propagation speed was estimated by dividing v.r. electrode distances by the propagation delay. For calculating the rhythmicity index for swimming and struggling, we followed the method in Li et al., 2010 ^52^. The v.r. recordings were first rectified after manually truncating stimulating artifacts to 0 µV. Then an event-triggering threshold was set at ∼ 5 times standard deviation obtained from traces clear of fictive swimming or struggling bursts. After generating the distribution histogram of event intervals, the first trough and peak event counts were used to calculate rhythmicity, i.e. the difference between peak and trough counts divided by their sum. For estimating the phase distribution of motoneuron spiking within struggling cycles, spiking events were identified by threshold crossing. Rostrocaudal phase delay then was calibrated following Green and Soffe ^28^: Standard phase delay = 0.185 - 0.00225 x cycle period, over a distance of 1.65mm between two longitudinal recording locations. If the recorded motoneuron was L mm rostral to the v.r. recording position, then the calibrated phase delay was standard phase delay multiplying L/1.65. If the motoneuron was recorded from the side opposite to the v.r. recording, then 0.5 was added to the calibrated phase delay.

### Immunohistochemistry

Tadpoles at stages 37-39 were fixed in 4% paraformaldehyde (PFA) on ice for 1.5 h and then rinsed in phosphate-buffered saline (PBS) before being dehydrated in methanol and bleached in a solution of 10% hydrogen peroxide in methanol overnight to bleach of the black pigmented cells. Then tadpoles were rinsed with pure methanol to remove hydrogen peroxide, rehydrated and dissected to remove the yolk and most part of the ventral body, before undergoing the CUBIC clearing protocol ^38^. CUBIC-L solution contained 10% Triton X-100 and 10% N-butyldiethanolamine in distilled water. CUBIC-R solution included 10% urea, 5% N,N,N’,N’-tetrakis, 10% Triton X-100 and 25mM NaCl in distilled water. CUBIC Refractive Index (RI) matching solution included 45% antipyrine and 30% nicotinamide in distilled water. Dissected tadpoles were first immersed in 50% CUBIC L+R in distilled water for 4 hours and 10 minutes at room temperature (RT) on a shaker, and then for 40 more minutes in 100% CUBIC L+R on a shaker at 37°C. After six six-minute washes in PBS, tadpoles were transferred in 10% normal donkey serum (Jackson ImmunoResearch, US) in 0.3 % PBS-Triton X-100 (PBST) for one hour at room temperature on a nutator, and then incubated with primary antibody for 4 days at 4°C on a shaker. At the end of antibody incubation, tadpoles were washed in PBST five times for one hour each, and transferred in the secondary antibody for one day at 4°C on a shaker. After 5 more washes in PBST of one hour each, tadpoles were left soaking in CUBIC RI matching solution overnight, before being mounted in homemade cavity slides in CUBIC RI matching solution. The primary antibodies used was goat anti-ChAT (1:200; Millipore). The secondary antibodies was a donkey anti-goat IgGs conjugated with Cy3 (1:500; Life Technologies).

### Image acquisition and cell count

Whole-mount preparations have been imaged using an Andor Benchtop Confocal-BC43 equipped with a 561 nm laser and Fusion software (Oxford Instruments, UK). Multi-image confocal stacks have been acquired with a 20x air objective (1 µm z steps), and a 10x air objective (1.8 µm z steps). Images were processed using Imaris (Oxford Instruments, UK) and Fiji in order to remove unnecessary z layers and add annotations in 3D-tiled imaging files. Automated cell counts based on machine learning algorithms in either Imaris or Fiji proved unreliable. Therefore, ChAT-positive neurons were annotated manually by means of “Spots and Surface” function in the Imaris software. In both 3D and slice “Spots and Surface” environment, clear objects with spheric and ovoidal shapes, often adjoining axon-like processes, were marked as neurons. As specimens were rarely mounted in perfect dorsal views, normally only one side of the CNS with better view of the spinal cord was used for cell counts. ChAT-positive cells rostral to the first myotome, presumably belonging to cranial motor nuclei, were excluded. Two additional reference markers were added to the rostral and caudal tadpole extremities and a third was added at the midbrain/hindbrain border. The coordinates for the reference markers were used to extrapolate the longitudinal locations of ChAT-positive cells.

### VT modelling

The VT movement simulations were carried out and visualised in our custom-written software Sibernetic-VT ^36^, which was originally developed to simulate the biomechanics of C. elegans movement ^53–55^. The interaction between objects was based on the Predictor-Corrector Incompressible Smoothed Particle Hydrodynamics algorithm (PCISPH)^56^. The reconstruction of VT was based on a series of cross sections of a real tadpole at different longitudinal positions, with numbers and shapes of VT muscle segments resembling the structure of a real tadpole. A particle-spring model was used, in which tissue density was represented with particle mass and the properties of springs connecting neighbouring particles represent tissue stiffness. The medium in which VT is placed is incompressible liquid that imitates water density, viscosity and surface tension at 20°C. The tadpole body was made of 14,141 particles whereas the virtual tank contained 33,684,488 water particles. The dimensions of the virtual water tank were 17.8 cm x 8.8 cm x 3.6 cm. The virtual tank wall was constructed from 187,000 stationary particles and the liquid particles did not penetrate through the wall. The VT belly has a density of 1.060 g/cm^3^, higher than the density of 1.035 g/cm^3^ in the notochord, muscles and other body tissue. The overall average tadpole tissue density is 1.04 g/cm^3^. As for real tadpoles, the VT notochord is the most rigid part of the body. Muscle particles are connected with contractile springs to simulate the contraction of “muscle fibers” (14 springs at muscle segments 1-8, 21 at segments 9-16 and 7 at the remaining segments). Adult frog muscles consisting mostly of fast twitch fibers have an average twitch stress at about 100 kN/m^2^ ^57,58^. *Xenopus laevis* tadpoles have superficial slow-twitch muscle fibres and deep fast-twitch fibres surrounding the notochord ^59^. Without knowing the extent of muscle activation, we gave a maximal contraction force of 3.3 µN for each single spring with an average of 30% activation. This gave the VT a range of body bends similar to that seen in real tadpole swimming and struggling.

These contractile springs are activated by 311 motoneurons on each side of the VT spinal cord and hindbrain (see below). All particles have a mass of 10 µg and are also subject to gravity with g = 9.8 m/s^2^. Distances, velocities, accelerations, time, force, mass, density and pressure were calculated in standard international units ^36^. The time interval for the numerical integration of equations was 0.01 ms. The videos generated from the simulations had frame rates of at least 1000 fps.

A pair of virtual forceps comprising 45, 994 particles were used to grip the VT’s head. The grip strength and friction were simulated by narrowing the gap between forceps tips from 100% of the tadpole head width to various degrees. Based on Hooke’s law, the compression force (N) on an elastic material is equal to Young’s modulus of elasticity (E) multiplied by the area (A) and deformation (Δw): N=E*A*Δw. The tadpole skin only contains two layers of cells and all parts inside the tadpole lack connective tissue. We hence assume standard E to be low at ∼700 Pa, comparable to brain tissue ^60^. Approximating the VT head as a sphere, the contact area (A, in mm^2^) between the forceps and each side of the head increases with deformation (Δw): A = 1.35Δw - 0.0043 mm^2^. The friction force F (in µN) resulting from head compression is given by

F = 2*μ*N*Δw = 2*μ*E*(1.35Δw - 0.0043)*Δw = 2*0.08* 700 *(1.35Δw - 0.0043)*Δw = 151.2Δw^2^ - 0.48Δw

where μ is the friction coefficient between the tadpole skin and the stainless steel forceps (Supplementary Fig.1). It was estimated by placing anaesthetised tadpoles on the surface of forceps. The forceps were tipped by a slowly increasing angle (α) until the tadpole started to slide down. At the threshold angle for the tadpole to slide, the friction is close to the gravity pull. Therefore, the friction coefficient (μ) could be calculated as: μ = tan α. The threshold slide angle was 4.5°, giving a friction coefficient of around 0.08 (Supplementary Fig.1C). The virtual forceps with smooth inside surface when they compress the VT head by 0.22 times h.w. could not hold the tadpole weight. Seven small ridges 52 µm in width evenly separated by 52 µm were therefore added to the end of the virtual forceps inside the surface on both tips to simulate the uneven surface of real forceps and increase the friction. The actual friction was measured by increasing a pulling force to the VT gradually until VT breaks free from the forceps. For compression by 0.22 times h.w., the friction was 6.8 µN, equivalent to 12.16 times the VT weight in water. After similar tests were carried out at different compression levels, an equation was fitted to capture the relation between VT head compression amount (C) and friction between VT and the virtual forceps (F): F = 49 Δw ^2^ + 19.8 Δw (Supplementary Fig.1B_2_). Theoretical calculations of friction based on Hooke’s Law and the friction tests on VT produced results in comparable ranges with some discrepancies, which may have resulted from imprecise contact area estimations.

In order to drive the movement in virtual tadpoles (VT), we generated artificial bursting activity of motor neurons to match their firing patterns in whole-cell recordings during fictive struggling in immobilised tadpoles after extending motoneuron distribution in the spinal cord along the VT longitudinal body axis to the tip of the tail. In our previous studies ^36,40^, we considered a model with a spinal cord up to 2 mm long and motoneuron soma distribution was based on Roberts et al., 1999 (also see Fig.3A, dashed line). In this study, we extrapolated the distribution of motoneurons to extend their innervation of the VT myotomes to the full body length, which is 4.65 mm from the mid/hindbrain border. This involved adding extra motoneurons from the point ∼3 mm posterior to the mid/hindbrain border towards the tail, at a density decreasing from 4 to 2 motoneurons per 100 µm, resulting in a full length VT model with 42 myotomes and 311 motoneurons on each side.

To generate inter-spike intervals (ISI) in each struggling burst with a given number of spikes, three different approaches were tried: a) Experimental data of ISI for the rostral part (motoneurons less that 2.2 mm from head) and caudal part were used to construct cumulative distributions; In order to match the distribution of inter-spike intervals, we excluded intervals longer than 13 ms, which could potentially represent spikes from different struggling bursts. b) A shallow artificial neural network (multi-layer perceptron) was trained to produce the next ISI given the previous ISI as an input; c) A deep artificial neural network was trained to produce the next ISI using all previous ISIs as inputs. These methods produced bursts with similar statistical characteristics and resulted in similar movements of the VT.

### Pressure drag calculation

To determine if the moving tadpole is in laminar flow, we estimated its Reynolds number: Re = 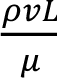, where *ρ* is the density of the fluid, *v* is the flow speed, *L* is a characteristic linear dimension and 𝜇 is the dynamic viscosity of the fluid. During struggling movements at ∼ 5Hz, if we consider that the tadpole tail passed describes a full circle and back, then *v* is around 85 mm/s and the distance covered *L* is 2 to 2.5mm. Considering water density at 1g/cm^3^ and viscosity of 0.00089 Pa·s at 20 to 25 °C, the formula gave us a Re of ∼ 190.6 for struggling. Given that Re >>1, we only considered the pressure drag ^61,62^ caused by the tadpole movement and given as 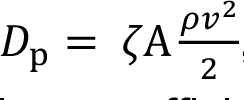, where *ρ* is the density of water, *v* is the flow speed, 𝜁 is the dimensionless coefficient of body shape resistance to flow (for a tadpole it is ∼ 1.18, assuming the tadpole coronal section is close to an ellipse), 𝐴 is the area of the body (for the whole tadpole it is ∼ 4.95 mm^2^).

To calculate the drag force created by each section of the tadpole (Fig.4A_1_), we calculated the area of each section separately assuming that the shape of the tadpole (side view) was approximately an ellipse. The formula for the upper/lower half of an ellipse is 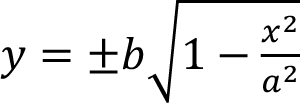 (where a and b are half the length of the long and short axes of the ellipse), which can be integrated along the body axis to calculate the area of each section assuming the centre of the ellipse is at the origin. The areas for the five sections on the rostral half of the tadpole are approximately as 0.64, 0.61, 0.55, 0.46 and 0.26 mm^2^. We assumed that the centre of mass was located at the middle of each section. Therefore, we calculated the velocity for the mass centre by averaging the velocity at the two ends of the section. We also treated each body section as a solid object without any elasticity, connected to its neighbors without any resistance. Pressure drag was applied to such solid sections in a direction opposite to the direction of their speed. This was projected to x, y and z axis. The total drag on each axis is the sum of the projected drag for all the segments. The calculations are valid when the velocity vector is perpendicular to the longitudinal axis of the body section. We simply calculated the drag by projecting the area of the section in the direction of velocity. Multiplying the area by the cosine of the acute angle between the body section and the velocity vector gave us the projected area. The vector in the direction of segment was obtained by subtracting the coordinates at the ends (Supplementary Fig.5). We provide drag and mass*acc calculation for VT videos, but not for real tadpole movement, because the former had higher frame rates and more accurate and reliable co-ordinates for all tracking points.

Statistical analyses were carried out using IBM SPSS22. Means were given with standard deviations (Mean ± SD) and compared using t-tests or one way ANOVA for normally distributed data. Otherwise, non-parametric independent-samples median test was employed. Pearson correlation was employed for normally distributed datasets and Spearman’s Rank correlation was used for non-normally distributed data.

## Acknowledgements

This research was supported by BBSRC (grant no. BB/T003146/1, BB/T002352/1, BB/T002549/1, BB/X005038/1). We thank Dr Lora Sweeney for helping with the immunostaining and Prof Alan Roberts, Drs Steve Soffe and Maarten Zwart for commenting on the manuscripts.

---

**Supplementary Fig. 1.**
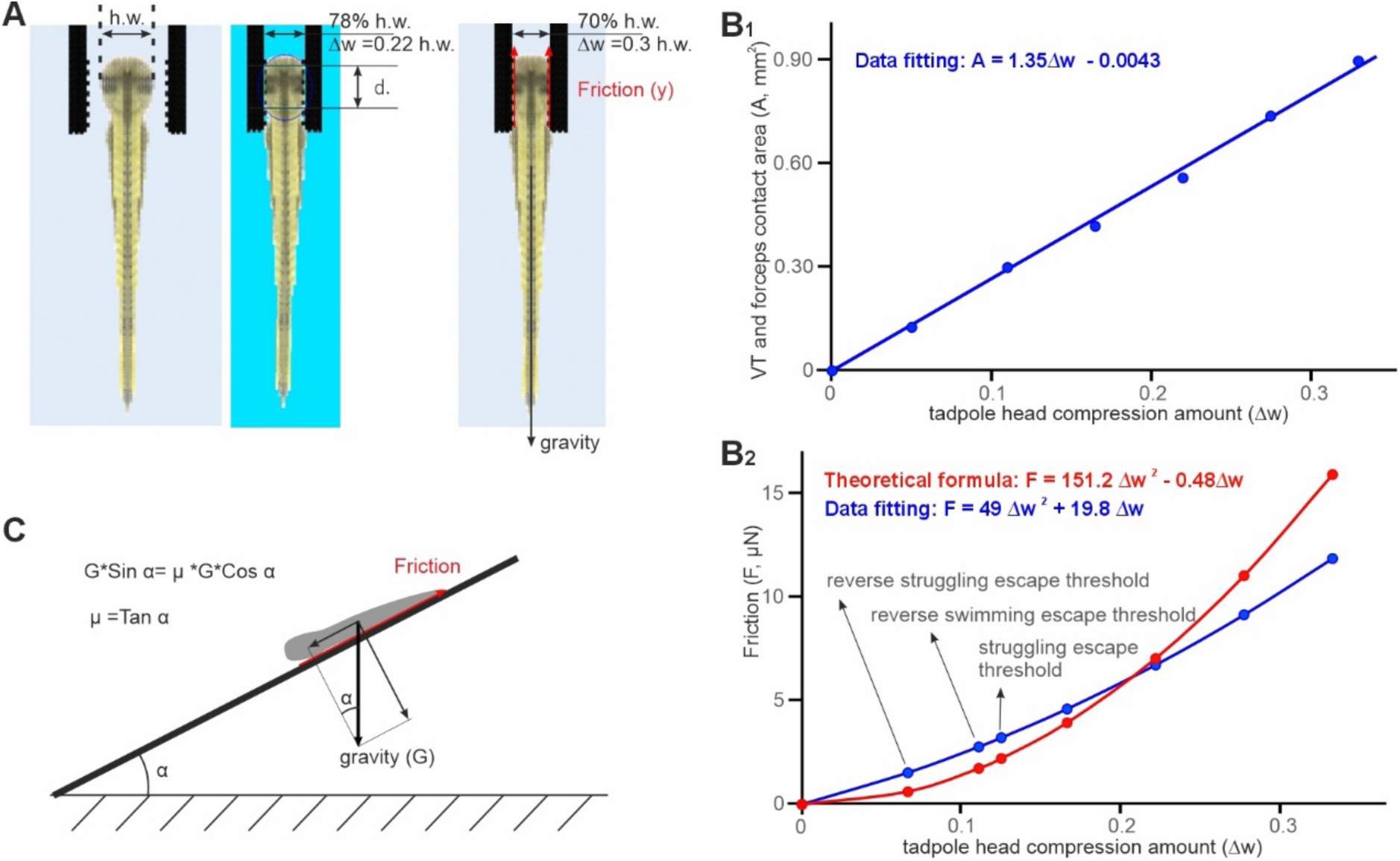
*Modelling friction between the VT and a pair of gripping forceps. **A**. Diagram showing the shape of virtual forceps with micro ridges at the tip to increase friction. h.w. is the VT head width (0.83mm, 100%). d. is the diameter of the circle of contact area. Red arrows indicate direction of friction, which is opposite to the direction of pull, i.e. gravity. **B******1****. The relation between amount of VT head compression and total contact area (A) between the virtual forceps inside surface on each side and the VT: A =1.35* Δw *-0.0043. Head compression (*Δw*) is converted to proportion of tadpole head width, i.e. compressed amount divided by h.w. B2. The relation between Friction (F) and VT head compression. Red symbols, curve and equation is based on Hooke’s Law. Blue ones are based on series of simulations to measure friction followed by data-fitting. The escape friction thresholds for struggling, reverse swimming and reverse struggling are annotated. They show that struggling enables VT to escape from levels of head compression higher than reverse swimming and reverse struggling. C. Estimating friction coefficient µ between the stainless steel forceps surface (thick solid line) and the tadpole skin. At a tilting angle when the tadpole starts to slide, G*Sin α=µ*G* Cos α. Hence µ = Tan α*.

**Supplementary Fig. 2.**
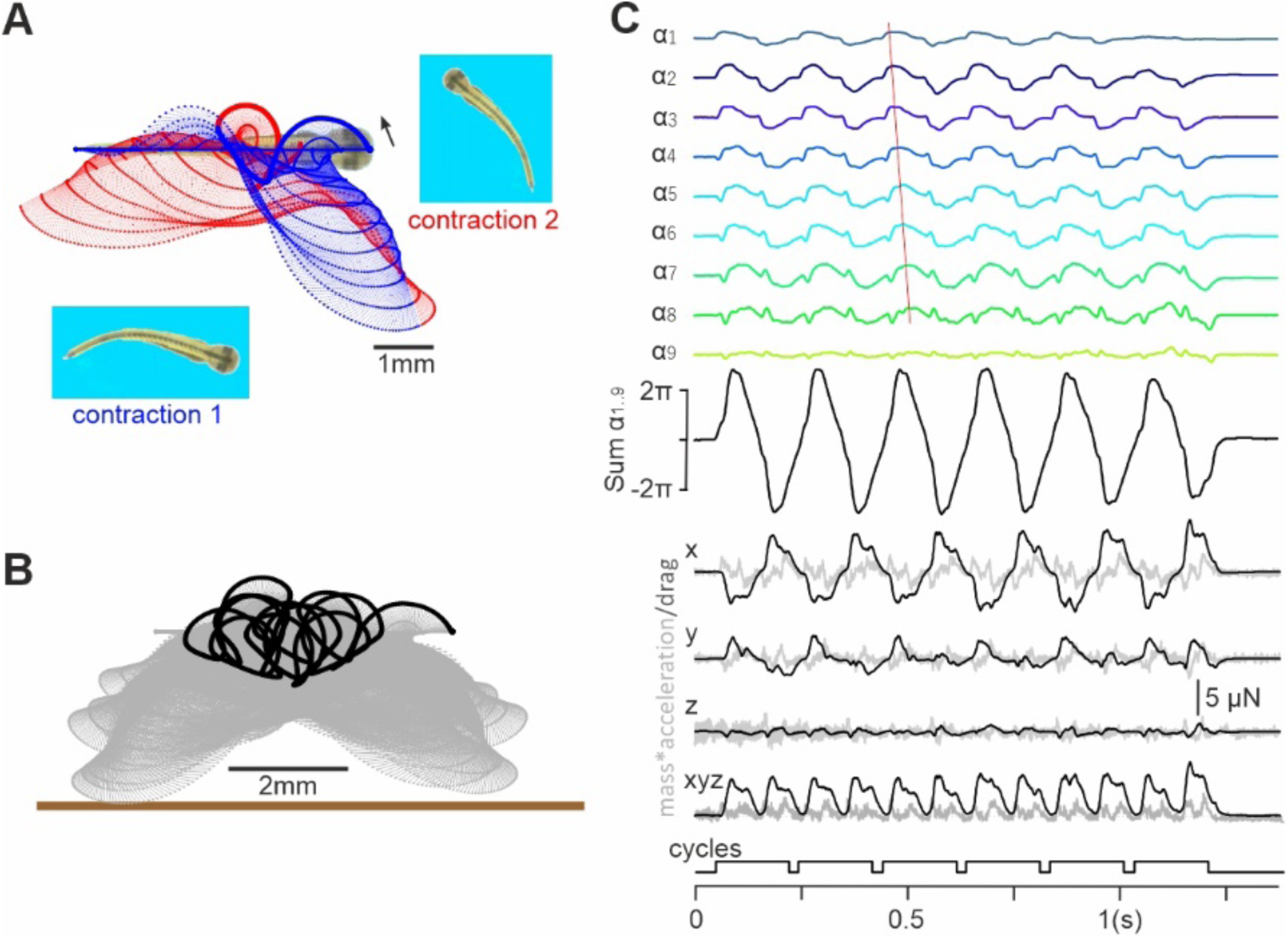
*Tracking free VT movement driven by reverse struggling commands. **A.** The movement trajectories of p1-11 along VT body axis of the first (blue symbols, lines) and second (red symbols, lines) contractions. Screenshots show VT body curvatures at the end of contractions. Larger dots indicate p1 trajectories. **B**. VT movement trajectory for the total simulation time (grey dots and dashed lines). Large black dots are for p1. Thick brown bar indicate a virtual water tank wall. **C**.VT body curvature angles, mass*acc. (grey traces) and drag (black traces) in three axes. Surrogate motoneuron spiking data are from* Fig.4D.

**Supplementary Fig. 3.**
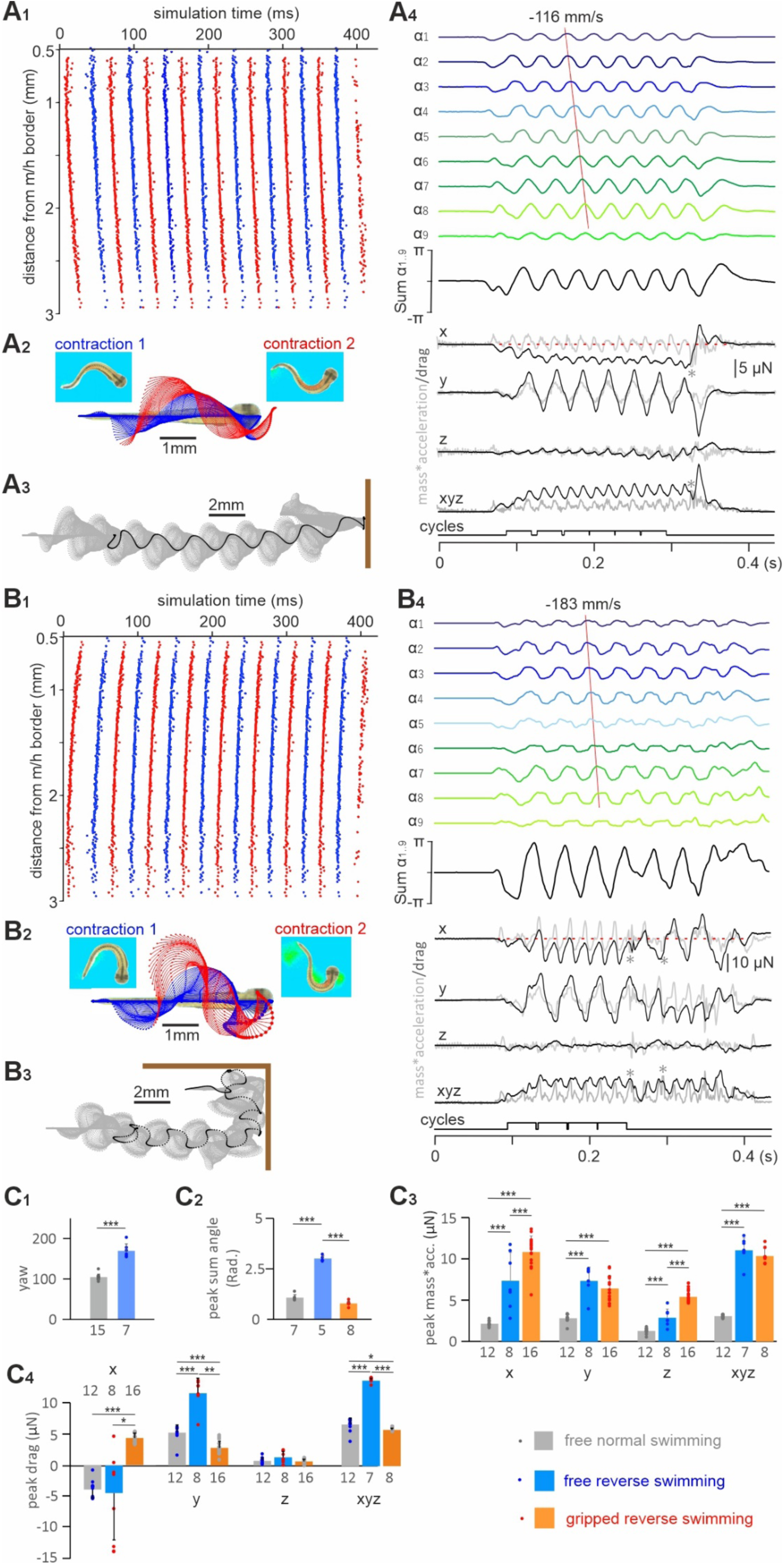
Simulating and tracking VT movement driven by normal and reverse swimming commands. **A**. Surrogate commands for normal VT swimming (**A1**) and its tracking analyses. **B**. Reverse, caudorostrally propagating swimming commands (**B1**) and its tracking analyses. **A2, B2**. Trajectories of 11 points along the VT tadpole body (on the xy plane) for the first (blue symbols, lines and text) and second (red symbols, lines and text) contractions. Screenshots show VT body curvature at the end of contractions. **A3, B3**. VT movement trajectory on the xy plane for the whole simulation time (grey dots and dashed lines). Large black dots indicate tracking point p1. Thick brown bars indicate virtual water tank walls. **A4, B4**.VT body curvature angles, mass*acceleration (grey traces) and drag (black traces) in x, y and z axes. xyz is the magnitude of 3D vector, combined from x, y and z. Cycles used for analyses are indicated. Asterisks indicates truncated mass*acceleration transients caused by VT touching the tank wall. Red dashed lines on x traces indicate 0. **C**. Comparing yaw of left-right movements driven by normal and reverse swimming commands (**C1**), peak of sum of angles (**C2**), peak mass*acc. (**C3**) and peak drag (**C4**) between free normal swimming, free reverse swimming and gripped reverse swimming (all pairwise two-tailed independent sample t-tests, *** indicates p < 0.001, ** indicates p < 0.01). Numerals beneath bars are numbers of contractions/cycles compared.

**Supplementary Fig. 4.**
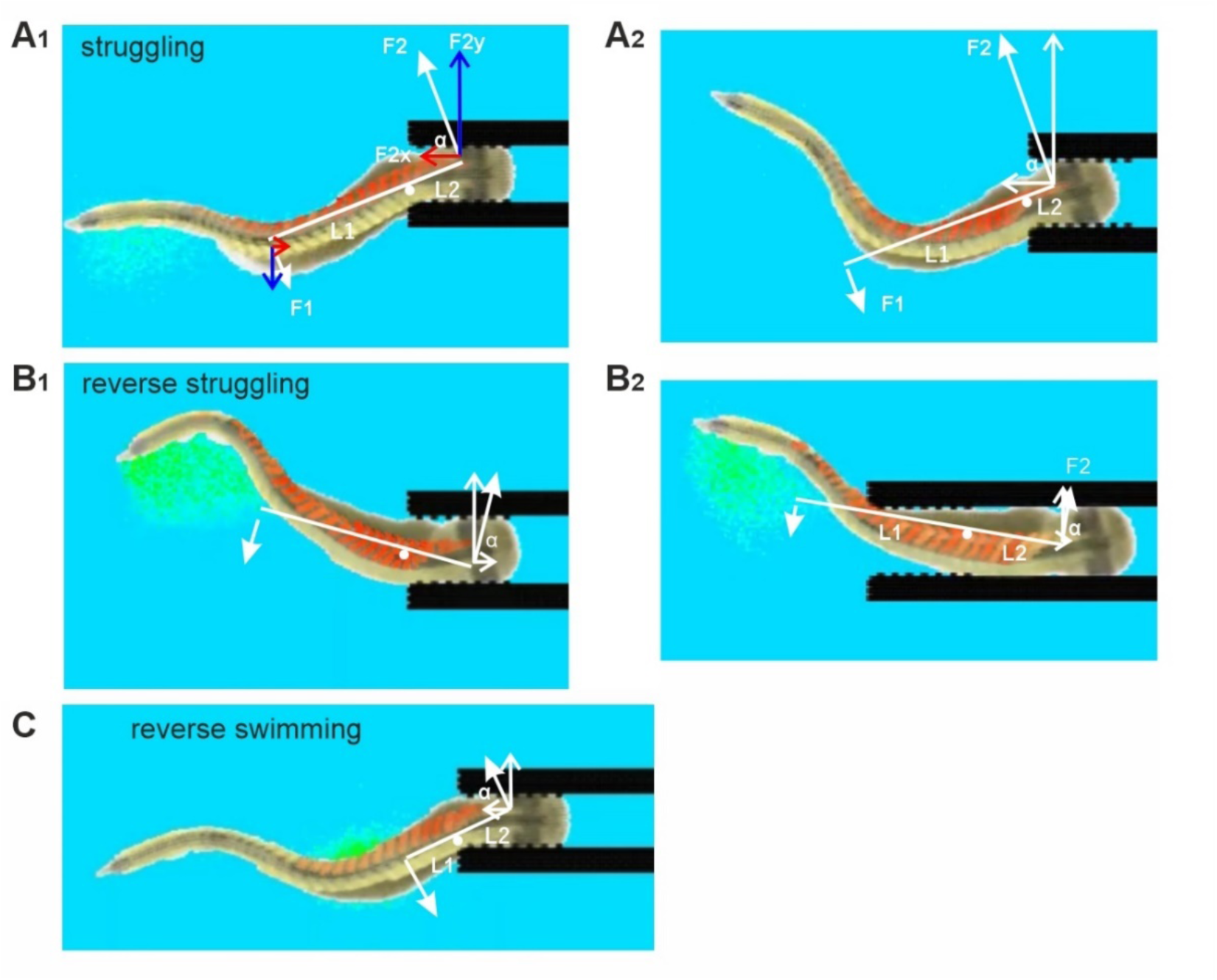
*Direct interaction between VT and the virtual forceps enables escape. The stiffened section of VT where muscles are contracted works like a lever with its contact point at the forceps tip as a fulcrum. The force F1 can be estimated as mass*acc. and results in a force F2 at the head: F2=F1*L1/L2, where L1 and L2 are the lengths of the lever ends. F2 has a horizontal axis (x) component that pushes the VT away from the forceps F2x= F2*cos(α), where α is the angle between F2 and the x axis. **A******1****. Struggling movement generates F2x that overcomes the friction between VT and the forceps and helps the VT to back out of the forceps. **A******2****. With less compression to the VT head, the ratio between L1 and L2 increases, generating larger F2x. Also the friction is smaller, leading to higher chance of VT backing out of the forceps. **B******1****. In reverse struggling, the body position when the muscles contract to produce a lever is opposite from struggling (c.f. **A******1****), generating a momentum that moves VT into the forceps. **B******2****. When VT is further into the forceps, the ratio between L1 and L2 decreases, while friction increases and the VT movements are more restricted, generating less force. This may explain why VT becomes stuck after moving deeper into the forceps. The fulcrum point is approximated. **C.** In reverse swimming, short swimming burst leads to less muscles contracting simultaneously so L1 is shorter than for struggling and consequently generates smaller F2x to overcome friction. All screen shots are taken when a maximal number of muscles in rostral VT are contracting simultaneously. The direction of F1 is determined by the direction of VT movement at the centre of mass of the moving segments (close to tracking position 4 in Fig.4A1)*.

**Supplementary Fig. 5.**
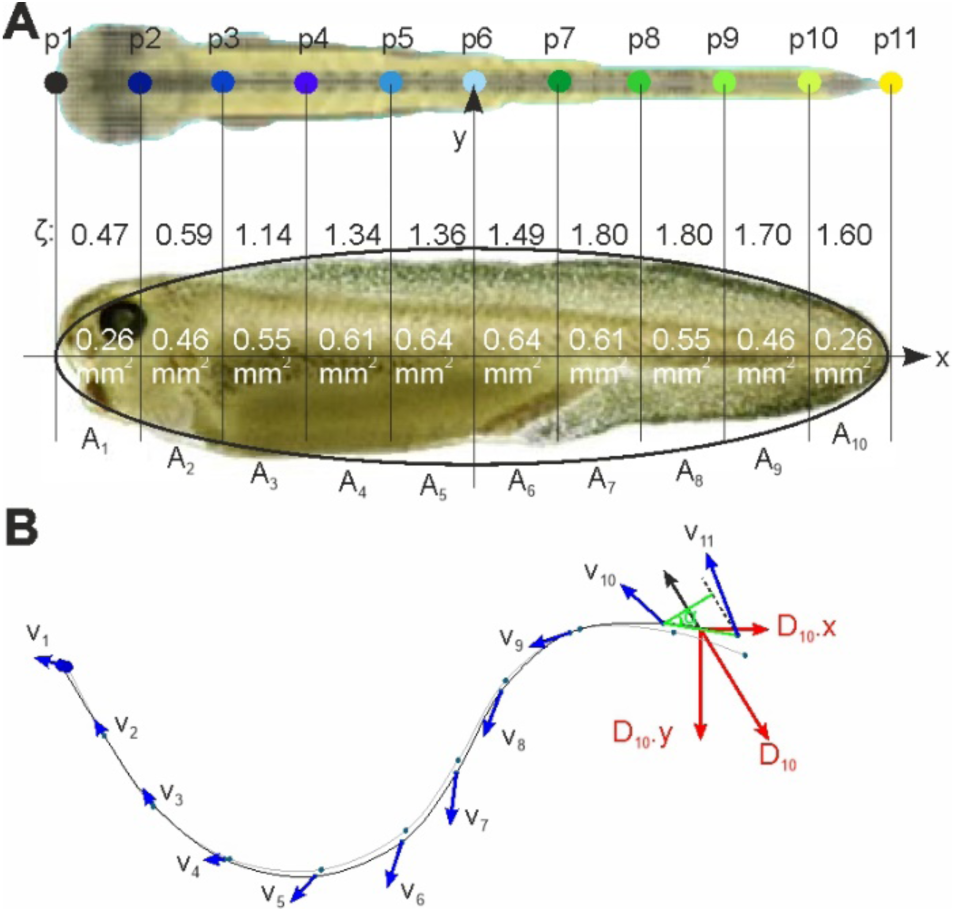
*Calculating pressure drag generated by VT movement. **A.** Estimating VT surface area. The side view outline of the VT could be approximated as an ellipse with a long axis of 5mm and a short axis of 1.25mm: y = ± 0.63*(1-0.16*x*^2^)^1^*^/^*^2^*. Half of the area for each of the 10 section can be calculated by integrating the formula between distances 0, 0.5, 1, 1.5, 2 and 2.5mm and resultant areas (A_1-10_) for each section are labelled. **B**. Diagram showing how to calculate and project drag onto the x and y axes using the section between tracking points p10 and p11, i.e. area A_10_. Thin lines and small dots are tracking positions of two consecutive frames in a video. Assuming the mass centre is at the middle point between p10 and p11, the mass centre velocity (black arrow) is calculated as the average of velocity vectors v_10_ and v_11_. Surface area (A_10_) is projected to the plane perpendicular to the averaged velocity vector: A_10_*cosine(α). The drag coefficient (ζ) for each section is estimated and given by the shape of its transection. Drag (D_10_) is calculated as: D_10_ = ζ_10_*A_10_* cosine(α)*ρv*^2^*/2, where ρ is the density of the water, v is averaged velocity, ζ is the coefficient of body shape resistance, which depends on the shape and cross-section area of the object against the water flow. Drag is then projected on axes x and y. Total drag on each axis is the summation of drags from all body sections, which are also used to calculate the total drag amplitude D_xyz_ = (D_x2_+D_y2_+D_z2_)_1/2_*.

